# A multi-kingdom genetic barcoding system for precise target clone isolation

**DOI:** 10.1101/2023.01.18.524633

**Authors:** Soh Ishiguro, Kana Ishida, Rina C. Sakata, Hideto Mori, Mamoru Takana, Samuel King, Omar Bashth, Minori Ichiraku, Nanami Masuyama, Ren Takimoto, Yusuke Kijima, Arman Adel, Hiromi Toyoshima, Motoaki Seki, Ju Hee Oh, Anne-Sophie Archambault, Keiji Nishida, Akihiko Kondo, Satoru Kuhara, Hiroyuki Aburatani, Ramon I. Klein Geltink, Yasuhiro Takashima, Nika Shakiba, Nozomu Yachie

## Abstract

Clonal heterogeneity underlies diverse biological processes, including cancer progression, cell differentiation, and microbial evolution. Cell tagging strategies with DNA barcodes have recently enabled analysis of clone size dynamics and clone-restricted transcriptomic landscapes of heterogeneous populations. However, isolating a target clone that displays a specific phenotype from a complex population remains challenging. Here, we present a new multi-kingdom genetic barcoding system, CloneSelect, in which a target cell clone can be triggered to express a reporter gene for isolation through barcode-specific CRISPR base editing. In CloneSelect, cells are first barcoded and propagated so their subpopulation can be subjected to a given experiment. A clone that shows a phenotype or genotype of interest at a given time can then be isolated from the initial or subsequent cell pools stored throughout the experimental timecourse. This novel CRISPR-barcode genetics platform provides many new ways of analyzing and manipulating mammalian, yeast, and bacterial systems.

**Teaser:** A multi-kingdom CRISPR-activatable barcoding system enables the precise isolation of target barcode-labeled clones from a complex cell population.

---

Cells are not homogeneous in any system. Although they proliferate and replicate the genome that encodes molecular regulatory programs into their progenies, they also change their statuses according to the dynamic response of gene expression patterns to environmental signals. As typically shown in multicellular organisms, cells self-organize through mutual molecular and mechanical communications and dynamically create complex structures. In these processes, spontaneous mutations in the genome sometimes impair the cellular program and cause cellular malfunction. Conversely, some other mutations confer growth or survival advantages to the cells, which can be beneficial or catastrophic to the system.

During cancer chemotherapy, resistant clones arise and expand with unique genetic, epigenetic, and cellular statuses, contributing to cancer recurrence and metastasis (*1–4*). In hematopoiesis, stem cells of an analytically indistinguishable group show fate-restricted differentiation patterns, in which some cells seem to be primed for specific lineages by incompletely understood factors (*5–8*). Similarly, *in vitro* stem cell differentiation and direct reprogramming experiments have demonstrated that “elite” clones reproducibly transform into target cell states (*9–11*). In laboratory microbial evolution experiments, different cells within the initial population dynamically expand and shrink their clone sizes through the acquisition of new mutations over multiple generations (*12–16*).

These views on clonal heterogeneity and cell lineage bias have been rapidly shaped by approaches that use DNA barcodes and deep sequencing for high-resolution clonal cell tagging. DNA barcoding involves integrating short and unique variable fragments of DNA—so-called “barcodes”—to individual cells by lentiviral transduction, transposon-mediated delivery, site-specific DNA recombination in the host genome, or plasmid transformation. In a given assay, the change in abundance of the barcoded clones is traced *en masse* by subsampling the cell populations and quantifying the DNA barcodes by deep sequencing read counts. Furthermore, the combination of barcode transcription and single-cell RNA sequencing (scRNA-seq) allows clonal lineages to be analyzed alongside cell states, revealing cell lineage-restricted state trajectories (*9, 17–23*). However, these measurements only provide snapshots of clone population sizes and gene expression statuses at the time of observation (*24*), limiting the analysis of the early causal molecular and/or environmental factors that derive specific fate outcomes of clones.

Did the chemotherapy-resistant clones exist in the initial cell population with the genetic mutations or altered cell state from the beginning? Did any molecular factor(s) underlie the observed stem cell differentiation fates? Was the progression of the specific clone conditional on the existence of any other clones? Flow cytometry cell sorting with immunostaining and emerging image cytometry cell sorting technologies enable the dissociation of heterogeneous cell populations into single cells with their observed phenotypes (*25–28*) but cannot do the same for a population of clones before they exhibit a phenotype of interest. Furthermore, any current high-content omic measurement requires the destruction of samples at the time of observation, which precludes the analysis of dynamic molecular and cellular behaviors within the same biological systems. The new concept of “retrospective clone isolation” has recently emerged to tackle the above-raised questions (*29–33*). In such a system, a barcoded cell population is first propagated, and its subpopulation is subjected to a given assay (Fig. 1A). After identifying a barcoded clone of interest, the same clone (or its close relative) is isolated in a barcode-specific manner from the initial or any other subpopulation stored during the experiment. The isolated live clone can then be subjected to any following experiments, including omics measurements and reconstitution of a synthetic cell population with the isolates.

**Fig. 1.**
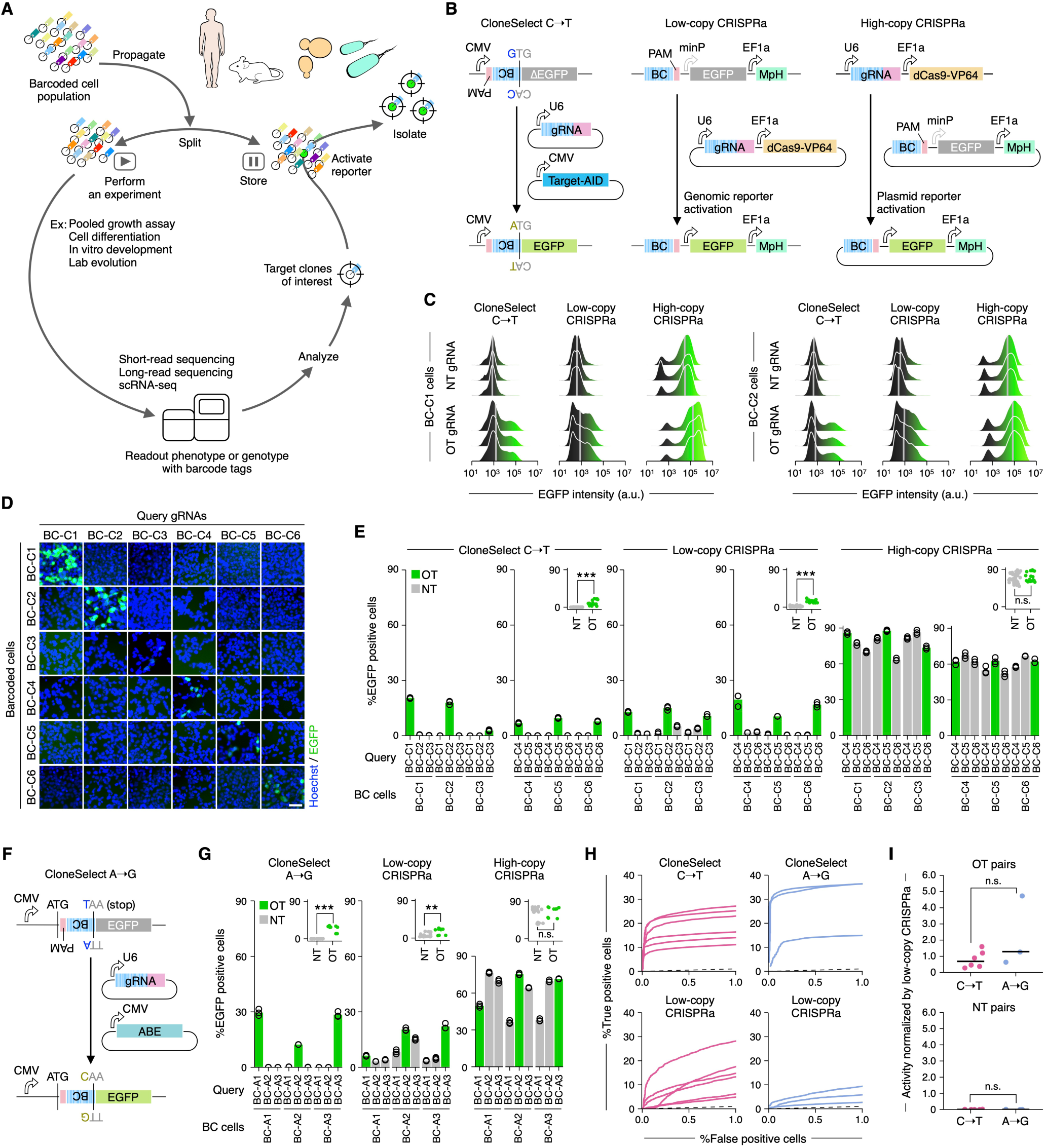
The CloneSelect circuit. (**A**) Conceptual diagram of retrospective clone isolation. (**B**) Different barcode-specific gRNA-dependent reporter activation circuits. CloneSelect C→T, low-copy CRISPRa, and high-copy CRISPRa. (**C**) Flow cytometry analysis of single-cell EGFP activation levels. (**D**) Barcode-dependent reporter activation of six barcoded cell lines by CloneSelect C→T. Scale bar, 50 µm. (**E**) Comparison of CloneSelect C→T, low-copy CRISPRa, and high-copy CRISPRa across the same barcode-gRNA pairs (n=3). For each approach, Welch’s t-test was performed to compare on-target (OT) and non-target (NT) activations. (**F**) CloneSelect A→G. (**G**) Comparison of CloneSelect A→G, low-copy CRISPRa, and high-copy CRISPRa across the same barcode-gRNA pairs (n=3). Welch’s t-test was performed to compare OT and NT activations. (**H**) ROC curves along varying EGFP intensity thresholds for target barcoded cells. Left, CloneSelect C→T and low-copy CRISPRa by the same targeting gRNAs. Right, CloneSelect A→G and low-copy CRISPRa for the same set of targeting gRNAs. (**I**) Performance comparison of CloneSelect C→T and CloneSelect A→G. Activated cell frequencies of OT and NT barcodes were normalized by activated cell frequencies of OT barcodes conferred by low-copy CRISPRa using the same targeting gRNA. The Mann-Whitney U test was performed to compare the two groups of datasets. \**P* < 0.05; \*\**P* < 0.01; \*\*\**P* < 0.001.

The idea of retrospective clone isolation has been implemented using CRISPR–Cas9 genome editing. In one form of the CRISPR activation (CRISPRa)-based approach (*29*), cells are tagged with DNA barcodes that are located upstream of a fluorescent reporter gene with a minimal promoter sequence. The barcoded cells are designed not to express the reporter gene before the guide RNA (gRNA)-induced transcription activation is triggered. Once a barcoded clone is identified for isolation, a gRNA targeting its barcode is introduced to the cell population with catalytically dead Cas9 (dCas9) fused to transcriptional activator(s) to trigger reporter expression of the target clone. This approach suffers from low specificity due to leaky expression of the reporter in the non-activated condition, presumably because of the effect of another constitutively active promoter neighboring the reporter. This effect can be mitigated by barcoding cells with a gRNA library pool and then labeling a target cell by delivering a plasmid that only encodes a reporter targeted by its corresponding gRNA, with no additional promoters present (*30, 31*). However, this strategy prohibits selection of the cells harboring the reporter plasmid using a secondary selectable marker. The CRISPRa-based approach requires continuous activation and maintenance of all components when a downstream experiment necessitates reporter expression in isolated cells. Furthermore, CRISPRa-based retrospective clone isolation has only been demonstrated in mammalian cell systems.

Genetic circuits based on DNA code alteration generally show highly specific input responses (*34*). A wild-type Cas9-based retrospective clone isolation has also been prototyped as a DNA code alteration-based approach (*32*). In this system, an out-of-frame start codon is followed by in-frame stop codons and a DNA barcode is encoded upstream of a reporter gene, whose translation is inactivated. The reporter translation can be restored by Cas9-induced double-stranded DNA break (DSB) and deletion through non-homologous end-joining (NHEJ) DNA repair at the barcode region, which stochastically removes the stop codons and brings the start codon into the coding frame. While this does not usually show unexpected reporter activation for non-target clones and confers permanent labeling of a target clone, the system’s sensitivity relies on stochastic DNA deletion, which can be a bottleneck to efficiency. Cas9-induced DSB is also cytotoxic and potentially impairs the target clone during the reporter activation procedure (*35, 36*).

Another approach has been proposed using RNA fluorescence *in situ* hybridization (FISH), where barcoded cells are first fixed, and a target transcribing barcoded clone can be labeled with a fluorophore probe and isolated by cell sorting (*37*). While RNA FISH is specific and sensitive, the isolated cells cannot be used for cell culture-based assays due to the fixation of the analyte.

Here, we report a new CRISPR base editing-based approach, CloneSelect, that overcomes the current technical constraints. CloneSelect is based on the restoration of reporter protein translation by base editing of an impaired start codon or an upstream stop codon in a barcode-specific manner. The new method is highly scalable, programmable, and compatible with scRNA-seq. Its specificity markedly surpasses other CRISPR-based systems. We also present the versatility of the method by implementing CloneSelect in human embryonic kidney (HEK) cells, mouse embryonic stem cells (mESCs), human pluripotent stem cells (hPSCs), yeast cells, and bacterial cells.

## Results

### Mammalian CloneSelect by C→T base editing

CRISPR base editing has widely been used to induce a single nucleotide substitution at a target genomic site without DSB (*38*). We first hypothesized that a C→T base editor (CBE)-based circuit (*39*) would enable highly sensitive, precise barcode-specific clone isolation with better performance than the previous CRISPR-based methods. In this CloneSelect C→T system, a barcode is encoded immediately upstream of a reporter gene whose start codon is mutated to GTG (Fig. 1B left). A constitutively active promoter transcribes the reporter-encoding region, but the impaired start codon makes its translation inactive. C→T base editing at the target barcode site enables the antisense strand of the first guanine (cytosine) to be substituted into thymine, restores ATG on the sense strand, and activates the reporter translation. To achieve this, we employed Target-AID, one of the CBEs with a narrow C→T base editing window in the gRNA target site (*36, 38*).

To examine the performance of CloneSelect C→T employing EGFP (enhanced green fluorescent protein) as a reporter, we compared it with two CRISPRa-based approaches. In the low-copy CRISPRa approach, cells are lentivirally labeled by barcoded CRISPRa reporters, each consisting of a barcode followed by a minimal promoter and EGFP with an MCP-p65-HSF1 (MpH) transcriptional activator expression unit encoded on the same vector (Fig. 1B middle). The reporter can be activated by introducing catalytically dead Cas9 (dCas9) fused to VP64 and a targeting gRNA fused to MS2 hairpins, interacting with the MCP (MS2 coat protein) domain of MpH to recruit transcriptional activators (*40*). In the high-copy CRISPRa approach, cells are labeled by gRNAs, in which gRNA spacers serve as barcodes (Fig. 1B right). Upon transfection of the high-copy CRISPRa reporter-encoding plasmids, the reporter expression can be induced in the cell encoding the targeting gRNA.

CloneSelect C→T was more sensitive than low-copy CRISPRa and more specific than high-copy CRISPRa in HEK293T cells. For each of the three approaches, we prepared two cell lines with the same set of barcodes (BC-C1 and BC-C2). CloneSelect C→T with on-target gRNAs activated the reporter expression in 19.17% (BC-C1) and 17.89% (BC-C2) of target cells while minimizing the activation by non-targeting gRNAs to less than 0.1% (Fig. 1C and Fig. S1A). Low-copy CRISPRa showed a lower reporter activation rate of 5.24–6.26% (Fig. 1C and Fig. S1B), and high-copy CRISPRa showed a very high background activation rate of 52.12–62.15% by non-targeting gRNAs (Fig. 1C and Fig. S1C). The reporter intensity of activated cells for CloneSelect C→T was slightly but significantly higher (1.76-fold) than that of low-copy CRISPRa (Fig. S1D). The sensitivity of CloneSelect C→T using Target-AID constructed based on a nickase Cas9 (nCas9) was higher than that constructed based on dCas9 (Fig. S1E). CloneSelect C→T was also demonstrated to activate the reporter expression in HeLa cells in a barcode-specific manner (Fig. S1F).

To systematically examine the orthogonality between barcodes and gRNAs across the three approaches, we expanded the analysis to six barcodes (Fig. 1D and Fig. S2A and B) and demonstrated that CloneSelect C→T and low-copy CRISPRa both specifically activated the reporter expression of target barcoded cells, where 2.81–20.46% and 10.39–19.53% of target cells were EGFP positive, respectively, with minimal non-target activation (Fig. 1E). Consistent with the preceding result, CloneSelect C→T exhibited significantly higher overall reporter expression level than low-copy CRISPRa (Fig. S2C) and the least false positive rate (Fig. S2D). High-copy CRISPRa showed substantially lower orthogonality (Fig. 1E and Fig. S2B and D), indicating that this approach is not practical for retrospective clone isolation. For CloneSelect C→T, we also optimized the input DNA amount without elevating the minimal false positive rate (Fig. S2E and F).

We also tested a wild-type CRISPR-based system in which the reporter expression interrupted by an upstream stop codon is removed by the barcode-specific DSB, followed by its stochastic deletion with NHEJ (Fig. S3A). The reporter activation rate of this approach remained at 2.14–2.30% for target barcoded cells with a relatively high-false positive rate of 0.51–0.56 % for non-target barcoded cells (Fig. S3B and C). Additionally, as an alternative to the EGFP reporter, we attempted to develop an mCherry reporter for CloneSelect C→T (Fig. S4A). However, the mutation of the start codon did not inactivate the fluorescent expression of the wild-type, presumably because the second methionine served as a start codon (Fig. S4B and C). We found that a stringent barcode-specific gRNA-dependent reporter activation can be established with the M1V (GTG)+M9A mutant without losing the fluorescence intensity level (Fig. S4D).

### Mammalian CloneSelect by A→G base editing

While exhibiting high efficiency and specificity in human cells, CloneSelect C→T is not applicable for clone isolation of bacterial species as they use GTG to initiate protein translation (*41*). We thus designed another system, CloneSelect A→G, using an adenine base editor ABE-7.10 that induces A→G base substitution at the gRNA target sequence (*42*). In CloneSelect A→G, following a constitutively active promoter and a start codon, a barcode encoding a TAA stop codon prevents downstream reporter translation (Fig. 1F). The stop codon can be removed in a gRNA-dependent manner by mutating the antisense strand of the thymine (adenine) to guanine, converting the stop codon into CAA (Proline). We compared the performance of CloneSelect A→G, low-copy CRISPRa, and high-copy CRISPRa (Fig. 1G and Fig. S5A–F) for three arbitrarily designed barcodes in HEK293T cells. Similar to CloneSelect C→T, CloneSelect A→G activated the reporter expression for 12.27–31.47% of on-target cells, whereas the non-target activation was maintained below 0.5% with high orthogonality of barcodes and gRNAs. Low-copy CRISPRa also exhibited barcode-specific reporter activation, but despite a lower reporter expression level, its overall false positive rate was higher than CloneSelect A→G (Fig. S5H). High-copy CRISPRa again showed a high background activation level.

To quantitatively compare the performances of CloneSelect C→T, CloneSelect A→G, low-copy CRISPRa, and high-copy CRISPRa, we analyzed reporter activation frequencies of on-target barcoded cells when a stringent false positive rate of 0.5% was permitted. CloneSelect C→T and CloneSelect A→G exhibited 10.05–24.88% and 14.12–35.19%, respectively (Fig. 1H). Low-copy CRISPRa examined with the same barcode sets also used for CloneSelect C→T and CloneSelect A→G exhibited 2.71–22.37% and 1.92–6.60%, respectively (Fig. 1H). In contrast, high-copy CRISPRa did not show reporter-activated cells at this threshold. The efficacy of the gRNA to recruit the effector Cas9, in general, has been known to depend highly on its targeting sequence (*43, 44*). To compare CloneSelect C→T and CloneSelect A→G, their cell activation frequencies for different target barcodes were normalized by those of low-copy CRISPRa for the same barcodes. As a result, we did not observe a marked difference in normalized performance between CloneSelect C→T and CloneSelect A→G (Fig. 1I).

### Isolation of barcoded human cells from a heterogeneous population

To examine if CloneSelect C→T can isolate target barcoded cells from a complex population, we next generated a barcoded lentiviral library by pooled ligation of barcoded EGFP fragments (Fig. S6A). The barcoded EGFP fragments were amplified by PCR using an upstream forward primer pool encoding semi-random barcodes and a common reverse primer (Fig. S6A and B). The barcode sequences were designed to be WSNS repeats (W=A or T; S=G or C) to avoid additional start codons from appearing. The ligation product was used to transform *Escherichia coli* (*E. coli*) cells, and transformant colonies on an agar plate were scraped to purify a lentiviral barcode library pool. We estimated that our protocol normally confers a barcode complexity of multiple hundreds of thousands (Fig. S6C) and confirmed most of them to have barcodes by DNA restriction digestion (Fig. S6D).

To perform a proof-of-principle demonstration, we transformed *E. coli* cells again with the constructed library pool, isolated barcoded plasmid clones into a 96-well plate, and pooled 93 that were confirmed to have single barcodes by Sanger sequencing (Fig. S6E). The plasmid mini-pool was used to transduce HEK293T cells with a low multiplicity of infection (MOI) of <0.1, ensuring a single barcode was introduced to each cell. We amplified the barcode region from the plasmid mini-pool and the transduced cells by PCR and analyzed them by high-throughput sequencing. We identified 115 barcodes (Fig. 2A) and found that the variation in barcode abundance was replicable in two independent sequencing library preparations (Fig. S6F) and largely inherited from that in the plasmid pool (Fig. S6G and H), suggesting no substantial barcode-dependent bias in lentiviral packaging and transduction.

**Fig. 2.**
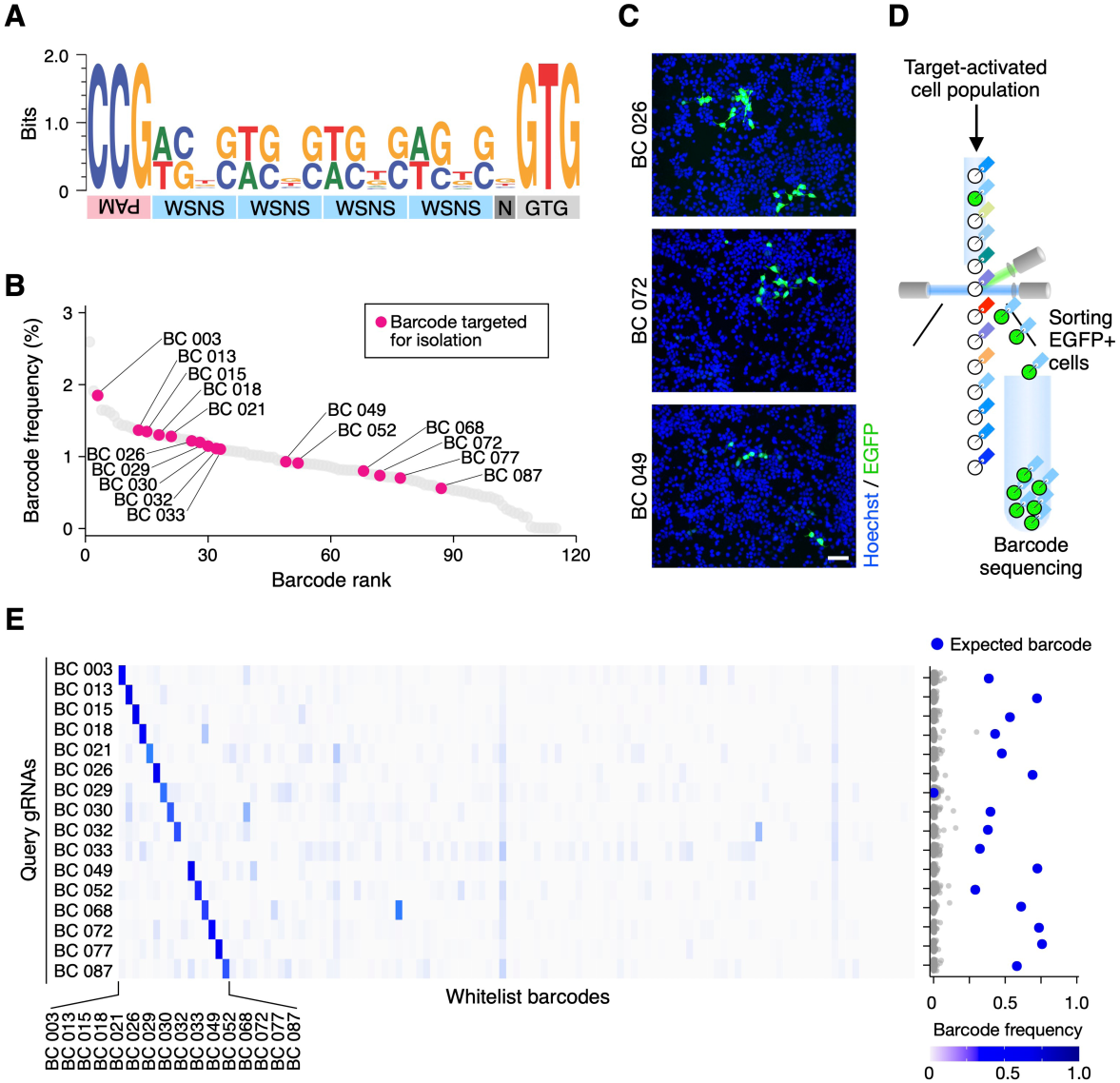
Barcode-dependent cell isolation from a HEK293T cell population. (**A**) Nucleotide compositions of barcodes in the mammalian CloneSelect C→T plasmid mini-pool. Five barcodes that had unexpected lengths were excluded from this visualization. The full barcode sequence list can be found in Table S1. (**B**) Barcode abundances in the cell population labeled by the mini-lentiviral barcode pool of CloneSelect C→T. (**C**) gRNA-dependent labeling of target barcoded cells in a population. (**D**) Flow cytometry cell sorting of reporter-activated cells. (**E**) Barcode enrichment analysis after cell sorting of the reporter-activated cells. Each row represents the barcode enrichment profile for each target isolation assay.

We then tested if we could enrich cells with 16 arbitrarily selected barcodes of different abundances in the cell population (Fig. 2B). For each target barcode, the cell population was co-transfected with the gRNA and Target-AID plasmids. After four days, we observed EGFP-positive cells in each assay (Fig. 2C) and sorted them by flow cytometry cell sorting (Fig. 2D). For each target barcode, its enrichment in the sorted cell population was analyzed by PCR and high-throughput sequencing (Fig. 2E). In sum, we succeeded in enriching the target barcoded cells in 15 out of 16 experiments (93.75%) with a relative barcode abundance of 29.18–75.75% after sorting. For the 15 successful targets, the mutated start codon was restored to ATG with an efficiency of 91.63–99.85% (Fig. S6I). A fraction of cells with the expected barcodes did not demonstrate the GTG→ATG mutation, suggesting that the cytidine deamination by Target-AID on the antisense strand might be sufficient to express the codon-repaired reporter transcripts.

### Isolation of clones identified in a scRNA-seq platform

To extend the utility of CloneSelect to isolate living clones identified according to their high-dimensional transcriptome profiles from a cell population, we made CloneSelect C→T compatible with 3′ capture scRNA-seq platforms and established scCloneSelect. In scCloneSelect, the barcode located upstream of the reporter with the mutated start codon (hereafter referred to as “uptag”) is paired with another barcode (“dntag”) downstream of the reporter followed by a hard-coded 30-nt poly(A) sequence (Fig. 3A). The dntag is captured as part of the short-read scRNA-seq reads through the common poly(A)-tailed 3′ enrichment strategy (*45, 46*) and used to refer its corresponding uptag for the reporter start codon restoration. This change in the circuit design did not affect the reporter activation performance of CloneSelect C→T in HEK293T cells (Fig. 3B and C and Fig. S7A and B) as well as the high orthogonality between barcodes and gRNAs (Fig. S7C and D). We also confirmed that dntag barcodes were transcribed (Fig. S7E) and efficiently captured by a scRNA-seq platform (Fig. S7F).

**Fig. 3.**
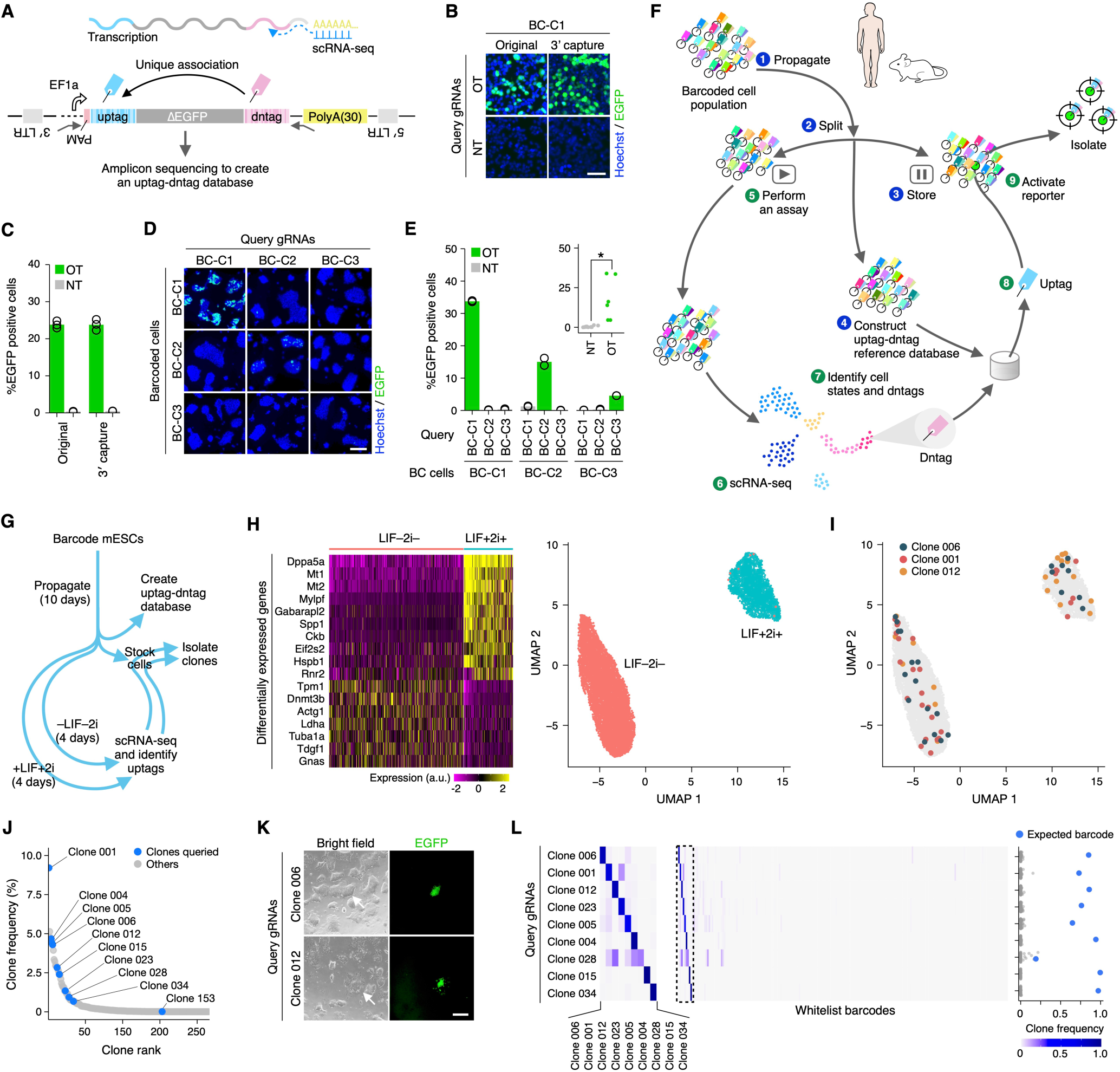
Isolation of live stem cell clones characterized by scRNA-seq. (**A**) scCloneSelect. (**B** and **C**) Barcode-specific gRNA-dependent reporter activation of the original CloneSelect C→T and scCloneSelect in HEK293T cells (n=3). Scale bar, 50 µm. (**D** and **E**) Barcode-specific gRNA-dependent reporter activation of three barcoded mESC lines by scCloneSelect. Target-AID was stably integrated prior to the barcoding. gRNAs were delivered by lentiviral transduction. Scale bar, 100 µm. Welch’s t-test was performed to compare on-target (OT) and non-target (NT) activations. \**P* < 0.05; \*\**P* < 0.01; \*\*\**P* < 0.001. (**F**) Schematic diagram of a scCloneSelect workflow to retrospectively isolate a cell clone demonstrating a gene expression profile of interest from a cell population stored before they demonstrate the target gene expression pattern. (**G**) mESC cell culture assays and clone isolation performed in this work. (**H**) scRNA-seq of mESC populations treated with LIF and 2i and those without LIF or 2i. (**I**) Distribution of cells for arbitrarily selected clones in the two-dimensional embedding of high-dimensional gene expression space by UMAP (uniform manifold approximation and projection). (**J**) Abundance of barcoded cell clones in the mESC population. The data was generated based on dntags identified by reamplifying the dntag reads from the original scRNA-seq libraries. (**K**) gRNA-specific activation of target barcoded clones in the mESC population. Scale bar, 50 µm. (**L**) Barcode enrichment analysis after cell sorting of the reporter-activated cells. Each row represents the barcode enrichment profile for each target isolation assay. The left heatmap was expanded from the dashed box area of the right heatmap.

One intriguing application of scCloneSelect would be to study the fate-determining factors of stem cell differentiation and cell reprogramming. scCloneSelect can be used to retrospectively isolate, from the initial population, cell clones whose states have been identified using scRNA-seq after differentiation. These clones could then be subjected to further analyses. We therefore tested if the scCloneSelect system is functional in selectively labeling target barcoded cells in mouse embryonic stem cell (mESC) and human pluripotent stem cell (hPSC) populations. In one of our approaches, a constitutively active Target-AID expression cassette was first stably integrated into a cell population by piggyBac transposon-based genomic integration, followed by lentiviral transduction of the cells with a scCloneSelect barcode library (Fig. S7G). The reporter activation was then triggered by introducing a targeting gRNA by lentiviral transduction or lipofectamine transfection. In mESCs, both of the gRNA reagent delivery methods activated the reporter expression in a barcode-dependent manner with no marked false positive activation, but lentiviral delivery of the gRNA showed an overall higher cell activation of 4.57–33.76% in comparison to 2.91–15.77% by the lipofectamine-based plasmid delivery method (Fig. 3D and E and Fig. S7H). The barcode-specific reporter activation was also successful in hPSCs, where targeting gRNA reagents were delivered by transfection (Fig. S7I). Additionally, we tested an approach that required a minimal number of steps, where hPSCs were first lentivirally barcoded, and the reporter was activated by electroporation with the targeting gRNA and Target-AID delivered together (Fig. S7J). This also enabled barcode-dependent reporter activation in hPSCs (Fig. S7K).

To demonstrate that barcoded cell clones identified with single-cell transcriptomes can be isolated from a barcoded cell pool sub-populated in parallel with the one used in scRNA-seq, we set up the following experiments using mESCs (Fig. 3F and Fig. S8A). In this experimental pipeline, a ΔEGFP (no start codon) fragment reporter is first amplified with forward primers encoding semi-random uptags of WSNS repeats accompanying a mutated start codon and reverse primers encoding random dntags. They are cloned into a common backbone plasmid *en masse* (Fig. S8B and C), and the constructed barcode plasmid pool is used to lentivirally barcode cells. Barcoded cells are then cultured to propagate cell clones (Step 1) and separated into three groups of sub-pools (Step 2). The first group is stored for later clone isolation (Step 3). The second group is used to identify the uptag-dntag combination reference database by PCR amplification and high-throughput sequencing (Step 4). The last group is subjected to a given assay, during which intermediate subpopulations can be stored at any point (Step 5). A cell population obtained during the assay is then analyzed by scRNA-seq to identify high-content gene expression profiles of single cells (Step 6). For a single cell demonstrating a cell state of interest, its dntags can be identified (Step 7), and the corresponding uptag can be retrieved from the uptag-dntag combination reference database (Step 8). Finally, using a gRNA targeting the identified uptag, the reporter expression of the clone containing the dntag is activated and isolated by cell sorting (Step 9). Because retroviral transduction, in general, is prone to having chimeric products of input vector sequences integrated into the genome through its high recombination activity (*47, 48*), the uptag-dntag database of the analyte population needs to be determined every time after the pooled transduction. We optimized the PCR protocol for the amplicon sequencing that identifies this database by minimizing chimeric PCR artifact products (Fig. S8D and E).

Using a plasmid pool with a barcode complexity of ∼150,000, we lentivirally transduced mESCs with Target-AID at an MOI of <0.1. Following the creation of a clone variation bottleneck by sparse sampling and 10 days of expansion, a subpopulation of the barcoded cells was used to construct the uptag-dntag database, in which 216 unique barcode pairs were identified (Fig. 3G). After preserving another subpopulation, the remaining cells were cultured with serum with LIF and 2i (LIF+2i+) or serum without LIF or 2i (LIF–2i–), to maintain or to lose pluripotency, respectively. Four days after, scRNA-seq was performed independently for the two conditions. While the RNA capture rates per cell of the two datasets were similar (Fig. S9A), the gene expression profiles of single cells were clustered into two distinct groups along with the culture conditions (Fig. 3H and Fig. S9B). Pluripotency marker genes, such as *Dppa5a*, were highly expressed in the pluripotency maintenance condition, whereas differentiation marker genes, such as *Tpm1*, were enriched in the non-pluripotent condition. Although the barcoded clones did not show a significantly biased distribution between the two conditions, we attempted to isolate the top 10 abundant clones in the scRNA-seq datasets (Fig. 3I and Fig. S9C and D) from the initial barcoded population. The abundances of these clones varied from 0.0133% to 9.21% in the initial population according to the analysis determined by the uptag-dntag database (Fig. 3J). Except for one experiment targeting Clone 153, we obtained a sufficient number of EGFP-positive cells after introducing the targeting gRNA using lentivirus followed by flow cytometry cell sorting (Fig. 3K and Fig. S9E). For each of the remaining nine clone isolation attempts, eight showed target clone enrichment with relative target abundances of 64.8–99.4%, whereas one (Clone 028) showed an enrichment degree of 18.9% (Fig. 3L). Furthermore, we isolated single cells from the EGFP-positive cell samples obtained for Clone 006 and Clone 012, expanded them (Fig. S9F), and confirmed the purification of the isolated clones (Fig. S9G). Accordingly, scCloneSelect was demonstrated to greatly enrich a target clone identified by scRNA-seq from a complex population with a success rate of 80% (eight out of 10).

### Yeast CloneSelect

Clonal barcoding approaches have also been used in microorganisms, such as yeast *Saccharomyces cerevisiae* (*S. cerevisiae*) and *E. coli*, to study their laboratory evolution and the genomic mutations accompanying clonal expansions of cells (*14, 16*). However, the analysis methods have been limited to time-course tracing of clone size dynamics. No technology has been developed to explore genetic, epigenetic, and other molecular factors contributing to laboratory evolution.

We next extended the CloneSelect C→T system for the yeast *S. cerevisiae* using mCherry as a fluorescent reporter (Fig. 4A). Like the mCherry reporter in mammalian cells, we also needed to truncate the first nine amino acids of mCherry to establish the reporter system (Fig. S10A). CBEs, including Target-AID, developed for mammalian species normally accompany uracil glycosylase inhibitor (UGI) to inhibit the base excision repair pathway, enhancing both the efficacy and purity of C→T substitution at the target site (*38*). However, Target-AID has been tested in yeast without UGI and was demonstrated to confer C→D (non-C) substitution at the target sequence at a high rate (*36*). We, therefore, constructed a yeast Target-AID with UGI and found that it did not largely impair the base editing activity (Fig. S10B–D) but as expected, greatly enhanced the frequency of resulting thymine at the target sequence (Fig. S10E). The efficient yeast CloneSelect C→T reporter activation was only possible by the addition of UGI (Fig. S10F). Similar to mammalian CloneSelect systems, yeast CloneSelect C→T was also demonstrated to activate barcoded cells in a highly target-specific manner with a sensitivity of 38.80– 44.13% by co-transformation of Target-AID and gRNA plasmids (Fig. 4B–D and Fig. S10G). Unlike mammalian cells, the labeled clones could be isolated by picking fluorescent colonies formed on a solid agar plate (Fig. 4E).

**Fig. 4.**
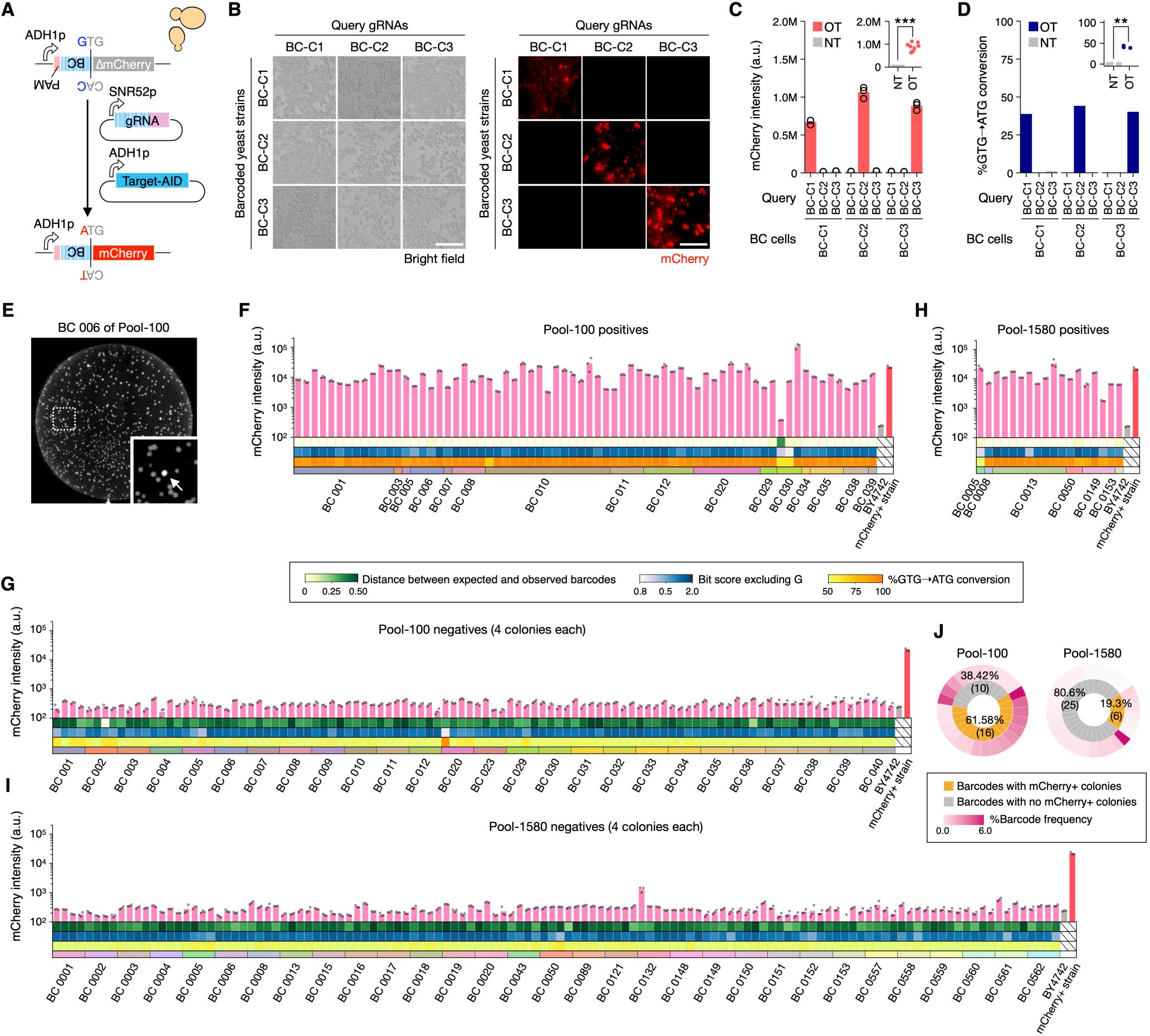
Yeast CloneSelect. (**A**) Yeast CloneSelect C→T circuit. (**B** and **C**) Barcode-specific gRNA-dependent reporter activation. Scale bar, 25 µm. Mean mCherry intensity measured by a plate reader was normalized by OD_595 nm_ (n=3). Welch’s t-test was performed to compare on-target (OT) and non-target (NT) activities. (**D**) GTG→ATG editing frequencies observed by high-throughput sequencing. Welch’s t-test was performed to compare OT and NT datasets. (**E**) Yeast colonies formed on a 10-cm agar plate after performing a target clone labeling in the yeast cell population of Pool-100. (**F–J**) Analysis of colonies isolated after clone labeling using each targeting gRNA. (**F**) mCherry positive isolates from Pool-100. (**G**) mCherry negative isolates from Pool-100. (**H**) mCherry positive isolates from Pool-1580. (**I**) mCherry negative isolates from Pool-1580. (**J**) Summary of the analysis results. \**P* < 0.05; \*\**P* < 0.01; \*\*\**P* < 0.001.

To test the sensitivity of the clone isolation strategy, we generated a barcode plasmid pool by a pooled ligation of semi-random barcode fragments to a backbone vector (Fig. S11A and B). Following a quality control of the pooled cloning reaction (Fig. S11C), *E. coli* cells were then transformed with the plasmid pool, and either 100 or ∼1,580 colonies were pooled to prepare barcode plasmid pools of defined complexities (hereafter referred to as Pool-100 and Pool-1580, respectively). After establishing the barcoded yeast cell populations of Pool-100 and Pool-1580 by yeast plasmid transformation, we arbitrarily targeted 26 and 31 barcodes of a range of abundances (0.83–5.79% for Pool-100 and 0.12– 0.37% for Pool-1580) to be isolated (Fig. S11D). For each barcoded clone isolation, its corresponding gRNA plasmid and Target-AID plasmid was co-transformed into the barcoded yeast cells, and fluorescent colonies were isolated, if any, together with four non-fluorescent colonies. The colony isolates were then cultured in liquid selective media in a microwell plate to measure the fluorescence intensities by a plate reader and barcode sequences were examined by PCR followed by Sanger sequencing (Fig. 4F–I). For Pool-100, 16 out of the 26 attempts (61.58%) conferred positive colonies, all of which, except for one of the three positive colonies obtained for BC 030, had the expected barcodes with a GTG→ATG conversion rate of 48.92–97.41% (Fig. 4J). For Pool-1580, six out of the 31 attempts (19.35%) conferred positive colonies, and all of them had the expected barcodes with a GTG→ATG conversion rate of 81.51–97.20%. The abundance of these successful barcodes in the Pool-100 and Pool-1580 yeast pools ranged from 0.120% and 5.78%, respectively, demonstrating that yeast CloneSelect can also isolate rare clones.

### Bacterial CloneSelect

Lastly, we also established a bacterial CloneSelect system for *E. coli* clone isolation. We first implemented CloneSelect A→G using an arabinose-inducible promoter for the EGFP reporter and an IPTG (Isopropyl β-d-1-thiogalactopyranoside)-inducible promoter for both the gRNA and ABE (Fig. 5A). The inducible promoter circuits showed an overall higher reporter expression in a barcode-specific manner (Fig. 5B). At the same time, the EGFP expression level by a targeting gRNA was higher than that by a non-targeting gRNA regardless of IPTG induction (Fig. 5C). The expected A→G substitution was also observed in a target barcode-dependent manner even for the no IPTG condition (Fig. 5D). These results collectively suggest that the background expression levels of the genome editing reagents with no inducer were sufficient to confer the base editing. Furthermore, when the EGFP positive control reporter was used, the arabinose-induced reporter intensity was suppressed by the targeting gRNA (Fig. 5E and Fig. S12A and B), suggesting a transcriptional silencing effect by ABE recruited at the target site.

**Fig. 5.**
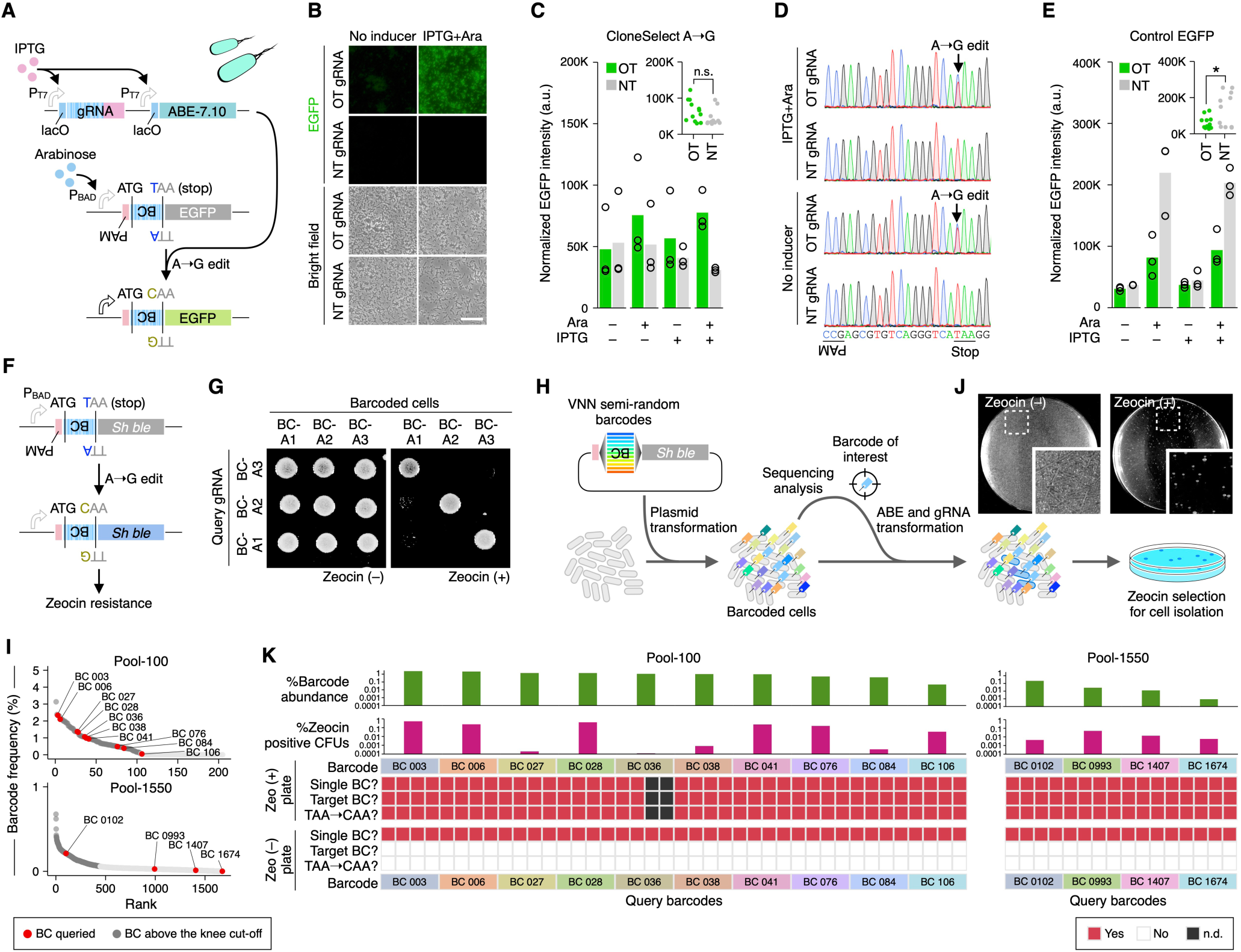
Bacterial CloneSelect. (**A**) Bacterial CloneSelect A→G circuit. ABE and gRNA expressions were controlled by IPTG-inducible promoters, and the EGFP reporter expression was controlled by an arabinose-inducible promoter. (**B** and **C**) EGFP reporter activation of *E. coli* cells under different inducer conditions. Scale bar, 25 µm. Mean EGFP intensity measured by a plate reader was normalized by OD_595 nm_ (n=3). Welch’s t-test was performed to compare on-target (OT) and non-target (NT) activities. (**D**) Base editing outcomes analyzed by Sanger sequencing. (**E**) Activities of the positive control EGFP reporter under the same conditions tested for (C) (n=3). Welch’s t-test was performed to compare OT and NT activities. (**F**) Zeocin resistance marker-based circuit. (**G**) Barcode-specific gRNA-dependent Zeocin resistance reporter activation. (**H**) Schematic diagram of a bacterial CloneSelect workflow using a drug selective condition for the target barcoded cell isolation. (**I**) Abundance of barcoded cells in Pool-100 and Pool-1550. (**J**) Colonies formed on Zeocin-selective and non-selective solid agar plates after performing the reporter activation of Clone 106 in the *E. coli* cell population of Pool-100. (**K**) Analysis of colonies isolated from Zeocin selective and non-selective plates obtained after clone labeling using each targeting gRNA. \**P* < 0.05; \*\**P* < 0.01; \*\*\**P* < 0.001.

Because the fold change in EGFP reporter intensity for the targeting gRNA compared to the non-targeting gRNA was not high (0.901–2.52 fold), we then sought to establish a drug-selectable system for bacterial CloneSelect. We first realized that when a Zeocin resistance gene (*Sh ble*) (*49*) was used for the barcode-specific reporter system (Fig. 5F), a constitutively active J23119 promoter conferred cell growth under the selective condition regardless of the upstream TAA stop codon (Fig. S12C). In contrast, the arabinose-inducible promoter was selective for the stop codon removal even without the arabinose supplement, suggesting that a small gene expression level is sufficient for drug resistance. We also found that targeting gRNA expression from the T7 promoter substantially dropped the number of colony-forming units (Fig. S12D), presumably because nickase Cas9 has known to be toxic to the bacterial cells (*50, 51*). Finally, we found that using the same setup as EGFP for the Zeocin resistance reporter with no inducer condition demonstrated cell growth in a gRNA-dependent manner (Fig. S12E). A Blasticidin S-resistance gene (*Bsr*) (*52*) also showed utility for gRNA-dependent cell growth with a similar experimental setup (Fig. S12F and G).

The Zeocin reporter system enabled the isolation of different barcoded strains with high specificity (Fig. 5G). To demonstrate barcoded cell isolation from a complex population, we constructed a pooled plasmid library for semi-random barcodes of VNN repeats (V=non-T), preventing the appearance of stop codons (Fig. S13A), and prepared sub-pool libraries by pooling 100 and ∼1,550 colonies, respectively (Fig. S13B; hereafter referred as to Pool-100 and Pool-1550). After establishing barcoded *E. coli* cell populations of Pool-100 and Pool-1550, we arbitrarily selected target barcodes and attempted to select cells having each of those barcodes by co-transforming the targeting gRNA and ABE, followed by Zeocin selection (Fig. 5H). We targeted 10 and four barcodes of a range of abundances in Pool-100 (0.047– 2.33%) and Pool-1550 (0.00089–0.211%), respectively (Fig. 5I). In every isolation experiment, the Zeocin selective conditions showed a substantially lower number of colonies than the non-selective conditions (Fig. 5J and K). For each of the successful target barcodes, except for BC 036 of Pool-100 in which we obtained only two colonies in the selective condition, four and four colonies were isolated from the selective and non-selective conditions, respectively, and their barcodes and base editing patterns were analyzed by Sanger sequencing. All isolates from the selective conditions had the expected barcodes and all of the isolates from the non-selective conditions had non-targeted barcodes. Accordingly, bacterial CloneSelect with the drug resistance marker system demonstrated great sensitivity and was able to isolate clones that were extremely low in frequency.

## Discussion

We describe a method, CloneSelect, which enables the isolation of target barcoded cells from a complex population using CRISPR base editing. Compared to the CRISPRa and wild-type Cas9-based systems tested in this study, CloneSelect demonstrated an ability to isolate cells with a higher sensitivity when a certain degree of false positive isolation was permitted. The low-copy CRISPRa experiments confirmed the reporter expression leak in similar existing methods (*29*). High background activity observed in the high-copy CRISPRa experiments supports the utility of a strategy implemented in the approaches COLBERT (*30*) and ClonMapper (*31*), whereby cells are labeled with gRNAs as barcodes, and target barcoded clones are activated by provision of a plasmid that encodes only the target gRNA-corresponding reporter with no constitutively active promoters. However, this approach of barcoding cells using gRNAs does not allow selected cells to maintain reporter activation for subsequent experiments, when necessary. CloneSelect largely benefits from the precision of base editing and the simplicity of altering the genetic code, while a CRISPRa-based system involves more additional endogenous factors to be coordinated for the target barcode-dependent reporter activation. Furthermore, engineering evolutionarily prevalent rules of the genetic code enabled us to implement the same idea across diverse species. We demonstrated the retrospective isolation of barcoded cells from complex yeast populations for the first time. While the isolation of barcoded *E. coli* cells has recently been demonstrated using barcode-specific CRISPR interference of a counter selection marker (*53*), we have shown that bacterial CloneSelect achieves the isolation of target barcoded cells at unprecedented sensitivity and selectivity.

When practically examining its ability to isolate target barcoded cells from complex populations, CloneSelect was able to retrieve rare, barcoded cells whose estimated abundances were as small as 0.558%, 0.013%, 0.120%, and 0.00089% in the human HEK293T, mouse ESC, yeast, and *E. coli* populations, respectively. The theoretical complexity of semi-random barcodes is limited to 4.19 x 10^6^ and 2.54 x 10^8^ for CloneSelect C→T and CloneSelect A→G, respectively. The sensitivities to obtain target barcoded cells that align with the same possible minimal abundance thresholds above are 93.75%,90.00%, 61.56%, and 100.00%, respectively. The limited sensitivity per isolation attempt could be explained by the general gRNA-dependent genome editing efficacy (*43*) since isolation success did not correlate well with the abundance of the target in a population. We suggest that the current sensitivity range of CloneSelect is sufficient in most prospective assays, with the expectation that multiple independent clones show a phenotype of interest that allows the user to have multiple attempts. More enhanced genetic circuits can be considered to improve sensitivity and specificity. We tested an OR-gate with C→T base editing of tandemly arrayed barcodes (Fig. S14A) and an AND-gate with A→G base editing of tandemly arrayed barcodes (Fig. S14B). The OR-gate did not function well and enabled an efficient reporter activation for only one of the three barcodes. The AND-gate enabled the three-input-dependent reporter activation, but exhibited a tight trade-off with the sensitivity, as expected. Given the high specificity of the system, introducing multiple independent CloneSelect barcodes per cell might be a solution to increase the sensitivity if necessary.

With CloneSelect, new wide-ranging experiments can be conceived in broad fields of biology. A CloneSelect-barcoded cell population can be subjected to any assay. Existing time-course scRNA-seq measurement strategies enable interrogation of different clonal lineages in a barcoded population alongside the dynamic changes in their gene expression landscapes, if the clone population sizes are not too small (*9*). In contrast, CloneSelect would allow the clones isolated from different time points from the progressing population to be analyzed by diverse approaches (Fig. 6A). Such non-transcriptomic analyses could include morphological analyses under a microscope and molecular analyses available for small amounts of input cells, but any currently available methods should be applicable for the downstream analysis if the isolated clones can be propagated for the given hypothesis.

**Fig. 6.**
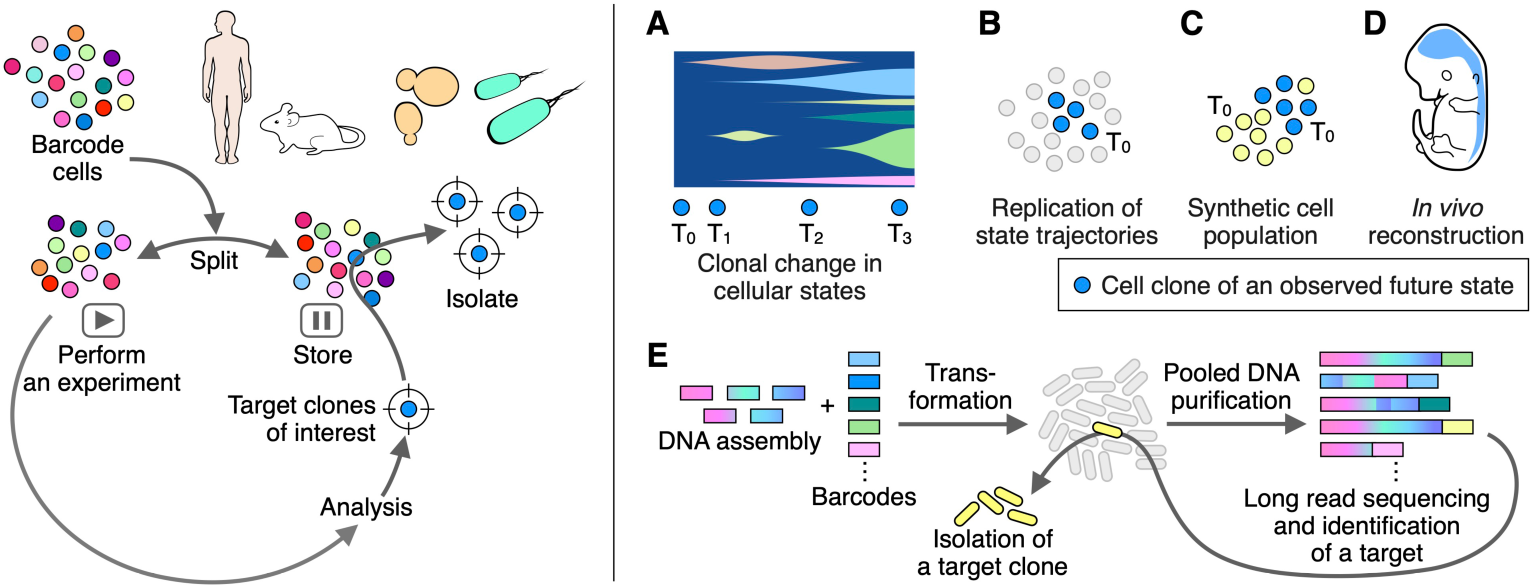
New biology directions made possible by CloneSelect. (**A**) Clonal analysis of molecular profiles in a complex population. (**B**) Replication of cell state trajectories. (**C**) Reconstitution of synthetic cell populations. (**D**) Transplantation of a fate-mapped clone. (**E**) New DNA assembly strategy.

Cells isolated by CloneSelect are alive. The clones isolated from the initial stage of a once-performed assay can be tested to see if they take the same developmental trajectories (Fig. 6B) or used to reconstitute a synthetic population with another cell population or other isolated clones (Fig. 6C). For example, a variety of hPSC lines have been reported to be favorable for different cell differentiation and organoid models (*54–58*), suggesting that there could also be fate priming of stem cell clones due to undiscovered intrinsic factors. CloneSelect enables mapping of cell states after differentiation and their subsequent isolation from the initial population. The fate-mapped stem cell clones could be used to engineer new stem cell-based models or high-quality stem cell therapeutics. Cell clones isolated from diverse systems can also be transplanted into animal models (Fig. 6D). Examples include xenotransplantation of a cancer stem cell clone and aggregation of a fate-mapped stem cell clone with an early embryo. As opposed to the CRISPRa-based system, the barcode-dependent reporter activation is irreversible. When a fluorescent reporter gene is used, the spatial distribution of the targeted clone and their interaction with others in a complex biological system can be traced.

In alignment with having DNA sequencing as a readout, CloneSelect would also promote the genetic engineering of cells and DNA assembly (*59–63*). Any cell engineering cannot be perfect—only a fraction of cells that go through an engineering process will harbor the target genetic product. Therefore, obtaining successful cells becomes difficult when their fractions in the reaction pool are very small. In such a task, through the barcoding and sequencing of the product cells, CloneSelect could retrieve target cells with the desired genetic engineering outcome. We also envision CloneSelect to largely improve DNA assembly in general (Fig. 6E). Currently, it is common practice to transform a DNA assembly reaction sample into *E. coli* cells in order to clone each assembly product, followed by colony isolation and sequencing-based confirmation. A pool of CloneSelect barcodes can be used to molecularly tag DNA assembly products. In this framework, the assembly products are used to transform *E. coli*, followed by the pooling of transformants and extraction of the pooled plasmid products from its subpopulation. The pooled plasmid product is then sequenced by long-read DNA sequencing to identify barcodes assigned to the desired product. Finally, the clone harboring the target product can be isolated using CloneSelect in a target barcode-specific manner. This strategy would enable the utilization of even inefficient DNA assembly protocols.

Accordingly, CloneSelect is a new way to precisely isolate target cell clones from complex populations. Its performance demonstrated in multi-kingdom species opens a vast array of possibilities to answer complex questions in diverse areas of biology.

## Acknowledgments

We thank the members of the Yachie lab at the University of British Columbia and the University of Tokyo, especially Sanchit Chopra, Sean Okawa, Sofia Romero, Yin Liu, Yeganeh Dorri, Madina Kagieva, and Rina Yogo, for their constructive feedback. We also thank Kaori Shiina, Shiro Fukuda, and Tara Stach for their technical support in high-throughput sequencing and Andy Johnson for the flow cytometry cell sorting. This study was funded by the New Energy and Industrial Technology Development Organization (NEDO), the Japan Agency for Medical Research and Development (AMED), the Japan Science and Technology Agency (JST), the Takeda Science Foundation, the SECOM Science and Research Foundation, the Canada Foundation for Innovation (CFI), the Canadian Institutes of Health Research (CIHR) CoVaRR-Net Project (all to N.Y.), and the Allen Institute Allen Distinguished Investigator Award (to N.Y. and N.S.). Authors were also supported by the JST PRESTO program (N.Y.), the Canada Research Chair program (N.Y.), the Ryoichi Sasakawa Young Leaders Fellowship (S.I.), the Taikichiro Mori Memorial Research Fund (S.I. H.M., and N.M.), the Japan Society for the Promotion of Science (JSPS) DC Fellowship (S.I., H.M. Y.K., and N.M.), the JSPS Overseas Research Fellowships (S.I.), Japan Student Services Organisation (R.S.), the Funai Overseas Scholarship (R.S.), the Takenaka Scholarship Foundation (R.T.), the Ministry of Science and Education (MEXT) Research Scholarship (A.A.), the Natural Sciences and Engineering Research Council of Canada (NSERC) Undergraduate Student Research Award (S.King), Michael Smith Health Research BC Scholar Award (R.I.K.G.), and the Michael Smith Health Research BC Trainee Award (A.-S.A). High-throughput sequencing data analysis was performed using the SHIROKANE Supercomputer at the University of Tokyo Human Genome Center.

## Author contributions

S.I. and N.Y. conceived the study. S.I., R.S., and M.T. performed all the major mammalian cell experiments. K.I. and M.T. performed all the yeast and bacterial experiments. O.B., N.S., M.I., and Y.T. performed a part of the experiments using mESCs and hPSCs. S.I., H.M., H.T., R.S., Y.K., and H.A. performed the high-throughput sequencing analyses. R.T., A.A., and N.M. performed preliminary assays and contributed to the design of the system. J.H.O., A.-S.A., and R.I.K.G. contributed to the optimization of the system. S.King supported the plasmid construction. K.N., A.K., and S.Kuhara also provided substantial ideas in designing the proposed system. M.S. supported the flow cytometry cell sorting. H.A. supported the high-throughput sequencing, the flow cytometry analysis, and the microscope imaging. S.I., R.S., K.I., and N.Y. wrote the manuscript.

## Materials and Methods

### Plasmids

Oligonucleotides were chemically synthesized by FASMAC, Integrated DNA Technologies, or Eurofins Genomics. All the oligonucleotides and cloning procedures used to construct the plasmids in this study can be found in Table S2. We used QUEEN version 1.2.0 (https://github.com/yachielab/QUEEN) to describe each plasmid construction and generate the annotated plasmid files in the GenBank (gbk) file format, embedding the full construction procedure (Table S2). A QUEEN-generated gbk file behaves as a quine code, where the QUEEN simulation code that generated the gbk file can perfectly be reconstructed from the file itself (*64*). We believe that providing the QUEEN-generated gbk files satisfies the mandate of reporting the reproducible protocols for the newly constructed plasmids. We also provided natural language descriptions for each plasmid construction step in the QUEEN code, allowing the user to retrieve the natural language description of materials and methods for each plasmid by simply executing “QUEEN --protocol_description --input [gbk file]” in a QUEEN-installed environment. A custom QUEEN wrapper that generated all QUEEN-generated gbk files can be found at https://github.com/yachielab/CloneSelect_v1/tree/main/QUEEN. We, therefore, do not provide the plasmid construction protocols here. All plasmid DNA sequences were confirmed by Sanger sequencing. The representative plasmids are in the process of being deposited to Addgene with their QUEEN-generated gbk files, which Addgene agreed to accommodate.

### Mammalian CloneSelect

#### Cell culture

##### HEK293T and HeLa cells

Human embryonic kidney 293Ta (HEK293Ta) and HEK293T Lenti-X cells were purchased from GeneCopoeia (#LT008) and Takara (#632180), respectively. Cells were cultured in Dulbecco’s Modified Eagle Medium (DMEM) (Sigma-Aldrich #11965084) supplemented with 10% Fetal bovine serum (FBS) (Gibco #16000044) and 1% Penicillin-Streptomycin (Wako #168-23191) at 37°C with 5% CO2 in a cell culture incubator. Cells were detached and passaged using 0.25 w/v% Trypsin-EDTA (Wako #203-20251) once the cells reached 70–90% confluency. The cell lines were regularly tested for mycoplasma contamination. When microscope imaging of HEK293T cells was performed with Hoechst 33342 (Invitrogen #H3570) counterstain, 100–200 µL of Collagen-I (Nippi #PSC-1-100-100) diluted in 5 mM acetic acid was added to each cell culture plate well, incubating for 30 min at 37°C. The collagen-coated plate wells were washed by 100–200 µL of 1x PBS before use.

##### mESCs

Under the approval by the Institutional Animal Care Committee of the University of Tokyo (RAC180003), mESCs were established from embryos of a 129(+Ter)/SvJcl (female) x C57BL/6NJcl (male) cross and maintained in Dulbecco’s Modified Eagle Medium - low glucose (Sigma-Aldrich #D6046-500ML) supplemented with 1% Penicillin-Streptomycin (Gibco #15140122), 1% MEM Non-essential Amino Acids (Wako #139-15651), 1% GlutaMAX Supplement (Gibco #35050061), 1% Sodium Pyruvate (Gibco #11360070), 15% FBS (Gibco #16000044), 100 µM/mL 2-Mercaptoethanol (Wako #131-14572), 10^3^ units/mL ESGRO Recombinant Mouse LIF Protein (Millipore #ESG1107), 3.0 µM CHIR99021 (GSK-3 inhibitor) (Wako #038-23101), and 1.0 µM PD0325901 (MEK inhibitor) (Toris #4423). A sufficient volume of 0.1% Gelatin (Sigma-Aldrich #G9391) in 1x Phosphate-buffered saline (PBS) (Takara #T9181 or Gibco #70011044) was added to each well such that the liquid could cover the entire surface and be aspirated after one hour at 37°C before plating cells. Cells were cultured in a cell culture incubator at 37°C with 5% CO2. The cell culture medium was replaced at least once every two days. The cell line was regularly tested for mycoplasma contamination.

##### hPSCs

CA1 human PSC (hPSC) line was used after approval by the Canadian Institutes of Health Research Stem Cell Oversight Committee. CA1 hPSCs were cultured using mTeSR Plus (STEMCELL Technologies #100-0276) cell culture medium in a humidified incubator at 37°C with 21% O2 and 5% CO2. Culture plates were coated with Geltrex LDEV-Free Reduced Growth Factor Basement Membrane Matrix (Gibco #A1413201). Dulbecco’s Modified Eagle Medium/Nutrient Mixture F-12 (DMEM/F12) (Gibco #11320033) was used to make a working solution of Geltrex (Gibco #A1413201) with 1:100 dilution. A sufficient volume of Geltrex was added to each well such that the liquid could cover the entire surface and be aspirated after one hour incubation at 37°C before plating cells. The cell culture medium was exchanged every other day after plating the cells. Regularly, cells were passaged as medium-sized clumps. Upon aspiration of the media, ReLeSR (STEMCELL Technologies #05872) was added, and cells were incubated at room temperature for about 1 min before the second aspiration. Cells were then placed inside the incubator for 4-5 min, added fresh media, and dissociated by pipetting up and down. Cells were then plated and placed inside the incubator. For single-cell passaging, we used TrypLE Express (Gibco #12604021). Cells were then placed inside the incubator for four min before adding a sufficient fresh media to stop the activity of TrypLE Express. Cells were then collected in centrifuge tubes, dissociated by pipetting up and down, and passed through a 40 µm cell strainer (Sarstedt #83.3945.040) to remove cell clumps. The tubes were then centrifuged at 300–400 g for 5 min and the supernatant was aspirated. The pellets were resuspended in a fresh media supplemented with 10 µM ROCK inhibitor Y-27632 (Tocris Bioscience #1254) for 24 hours to promote the survival of the single cells. For culturing H1 hPSCs, we used StemFit AK02N cell culture medium (REPROCELLAHS #RCAK02N) with Y-27632 (Cayman #10005583) added for one or two days after plating. Culture plates were coated with the recombinant Laminin-511 E8 fragment using iMatrix-511 Silk (MAX #892021). The cell lines were tested for mycoplasma contamination.

#### Mammalian CloneSelect barcode libraries

##### CloneSelect C→T barcode library

To generate the CloneSelect C→T barcode library, a semi-random oligo pool SI#679 encoding 5′-CCGWSNSWSNSWSNSWSNSNGTG-3′ was first chemically synthesized (Table S2), where the antisense strand sequence of the 5′-CGG-3′ PAM sequence and a quadruple repeat of WSNS (W=A or T; S=G or C) were followed by a mutated start codon (GTG). The WSNS repeat restricts additional start codons from appearing before the downstream reporter. An EGFP coding sequence was then amplified from pLV-eGFP (Addgene #36083) in 25 separate PCR reactions, each in 50 µL volume, composed of 1 ng/µL of pLV-eGFP template plasmid, 1.25 µL of 20 µM SI#679 oligo pool as a forward primer,1.25 µL of 20 µM SI#680 as a common reverse primer, 0.5 µL of Phusion High-fidelity DNA Polymerase (NEB #M0530), 10 µL of 5x Phusion HF Buffer (NEB #B0518S), and 5 µL of 2.5 mM dNTPs (Takara #4025), with the following thermal cycle condition: 98°C for 30 s, 30 cycles of 98°C for 10 s, 72°C for 10 s, and 72°C for 60 s, and then 72°C for 5 min for the final extension. The amplified fragment was digested by DpnI (NEB #R0176) for an hour at 37°C, pooled into a single tube (1,250 µL in total), and column purified using FastGene PCR/Gel Extraction Kit (Nippon Genetics #FG-91302). The purified fragment was then subjected to EcoRI-HF (NEB #R3101S) and XbaI (NEB #R0145S) digestion overnight at 37°C and purified again by FastGene PCR/Gel Extraction Kit (Nippon Genetics #FG-91302). To obtain a highly complex lentiviral plasmid pool, we performed a total of five ligation reactions using PCR strip tubes, each for a 50-µL reaction containing ∼30 fmol of EcoRI-XbaI digested pLVSIN-CMV-Pur backbone plasmid (Takara #6183), ∼300 fmol of the insert fragment, 2.5 µL of T4 DNA Ligase (NEB #M0202), and 5 µL of 10x T4 DNA Ligase Buffer (NEB #B0202). The reaction samples were incubated at room temperature for two hours and purified by FastGene PCR/Gel Extraction Kit (Nippon Genetics #FG-91302). The ligation sample was then used to transform NEB Stable Competent *E. coli* cells (NEB #C3040I). The transformation was performed in five reactions, each with 1,250 ng of the ligation sample transformed to 200 µL of the competent cells according to the manufacturer’s high-efficiency transformation protocol. Following one-hour outgrowth in SOC medium (NEB #B9020) at 37°C, cells were spun down and plated on a total of 25 LB agar plates containing 100 µg/mL Ampicillin (Wako #014-23302). After overnight incubation at 37°C, colonies formed on the plates were scraped by adding 1–2 mL ddH2O, pooled into a flask, and further incubated with 200–300 mL of LB liquid medium containing 100 µg/mL Ampicillin (Wako #014-23302) overnight at 37°C. We plated the transformation sample on agar plates with 500-fold dilution in triplicate and estimated the barcode complexity of the constructed library to be ∼6.8 x 10^5^. The plasmid library was finally purified by NucleoBond Midi-prep Kit (Macherey-Nagel #740410) and stored at –20°C before use. We performed the isolation of 16 random clones and triple restriction enzyme digestion using BsrGI-HF (NEB #R3575S), ClaI (NEB #R0197S), and PvuI-HF (NEB #R3150S), and confirmed that all the tested clones (16/16) contained the expected barcode inserts. To generate a low-complexity library for the proof-of-concept assays using HEK293T cells, we examined barcode sequences of 96 isolated clones by Sanger sequencing using a sequencing primer SI#471. After removing three clones observed to have mixed Sanger sequencing spectra in the barcode region, barcoded plasmids were pooled with an equimolar ratio and subjected to high-throughput sequencing and lentiviral packaging.

##### scCloneSelect barcode library

The scCloneSelect barcode library was prepared similarly to the CloneSelect C→T barcode library. An EGFP coding sequence was first amplified by PCR from pLV-CS-112 (Addgene #131127) using the semi-random oligo pool SI#679 as a forward primer and another oligo pool RS#244 as a reverse primer. The PCR was performed in 25 separate reactions, each in 40 µL reaction volume, composed of 0.12 µL of 10 ng/µL pLV-CS-112 template plasmid, 2 µL each of forward and reverse primers, 0.6 µL of Phusion High-fidelity DNA Polymerase (NEB M0530), 8 µL of 5x Phusion HF Buffer (NEB #B0518S), and 3.2 µL of 2.5 mM dNTPs (NEB #N0447) with the following thermal cycle condition: 98°C for 30 s, 30 cycles of 98°C for 10 s, 65°C for 10 s, and 72°C for 60 s, and then 72°C for 5 min for the final extension. The amplified barcode-EGFP fragment was pooled into a single tube (1,000 µL in total) and digested with 12.5 µL of DpnI (NEB #R0176) at 37°C for 1 hour. The digested product was then size-selected using FastGene PCR/Gel Extraction Kit (Nippon Genetics #FG-91302). The purified product was subjected to EcoRI-HF (NEB #R3101S) and XbaI (NEB #R0145) digestion overnight at 37°C and purified again by FastGene PCR/Gel Extraction Kit (Nippon Genetics #FG-91302). For backbone preparation, 25 µg of pRS193 lentiviral cloning backbone plasmid was digested by EcoRI-HF (NEB #R3101S) and XbaI (NEB #R0145) at 37°C overnight and size-selected using FastGene PCR/Gel Extraction Kit (Nippon Genetics #FG-91302). 1.25 µg of the digested backbone, 320 ng of the purified insert, 25 µL of T4 DNA Ligase (Nippon Gene #317-00406), and 25 µL of 10x T4 DNA Ligase Buffer (NEB #B0202) were mixed in a total volume of 250 µL and incubated overnight at 16°C. The ligation sample was then used to transform NEB Stable Competent *E. coli* cells (NEB # C3040I). The transformation was performed in a total of 17 reactions, each with 4 µL of the ligation sample transformed to 50 µL of the competent cells according to the manufacturer’s high-efficiency transformation protocol. Following one-hour outgrowth in SOC medium (NEB #B9020) at 37°C, cells were spun down and plated on a total of 15 LB agar plates containing 100 µg/mL Ampicillin (Wako #014-23302). After overnight incubation at 37°C, colonies formed on the solid agar plates were scraped by adding 1–2 mL ddH2O, pooled into a flask, and further incubated with 200–300 mL of LB liquid medium containing 100 µg/mL Ampicillin (Wako #014-23302) overnight at 37°C. We plated the transformation sample on agar plates with 300-fold dilution in duplicate and estimated the barcode complexity of the constructed library to be ∼1.5 x 10^5^. The plasmid library was finally purified by NucleoBond Midi-prep Kit (Macherey-Nagel #740410) and stored at –20°C before use. We performed the isolation of 20 random clones and confirmed the expected fragment insertion for 17/20 by genotyping PCR using the primer pair RS#147 and SI#514. From the same 20 clones, we selected six (including the three that did not yield the expected genotyping PCR bands), performed double digestion using EcoRI-HF (NEB #R3101S) and BamHI-HF (NEB #R3136S), and Sanger sequencing using sequencing primers SI#514 and RS#147 for uptag and dntag, respectively, and confirmed that all the tested clones contained the expected uptag and dntag inserts.

#### Lentiviral barcoding

##### Virus packaging

HEK293T cells were plated on either a 10-cm cell culture dish at a density of ∼2 x 10^6^ cells with 10 mL of the culture medium or 6-well cell culture plate wells at a density of ∼2 x 10^5^ cells/well with 2 mL of the culture medium one day before plasmid transfection. For packaging using a 10-cm cell culture dish, 3.0 µg transgene vector, 2.25 µg psPAX2 (Addgene #12260), 0.75 µg pMD2.G (Addgene #12259), and 18 µL of 1 mg/mL PEI MAX (Polysciences #24765-100) were dissolved in 1,000 µL of 1x PBS and applied to the cell culture. For packaging using a 6-well cell culture plate well, 489 ng transgene plasmid, 366.7 ng psPAX2, 122.3 ng pMD2.G, and 2.93 µL of 1 mg/mL PEI MAX (Polysciences #24765-100) were dissolved in 300 µL of 1x PBS and applied to the cell culture. The culture medium was changed to a fresh medium one day after transfection. The transfected cells were further incubated for 48–72 hours. The cell culture supernatant was then harvested and filtered with 0.22 µm pore size sterile syringe filters. The recombinant lentivirus sample was then aliquoted 500–1,000 µL each into 1.5-mL test tubes and stored at –80°C.

##### Virus concentration

To increase virus infection titer, we concentrated harvested virus samples using a polyethylene glycol (PEG)-based method (*65*) using PEG 6000 (Wako #169-09125) or Lenti-X Concentrator (Takara #631231). When PEG 6000 was used, ∼10 mL of recombinant virus sample was mixed with 2.55 mL of 50 w/v% PEG 6000, 1.085 mL of 4M NaCl, and 1.365 mL of 1x PBS in a 50 mL tube. The sample was continuously mixed using a rotator at 4°C for 90 min and centrifuged at 4,000 g and 4°C for 20 min. The supernatant was discarded, and the precipitated virus pellet was resuspended with 1.1 mL of Opti-MEM (Gibco #31985062) by pipetting and vortexing until fully dissolved, resulting in a 10-fold concentration of the virus sample. The virus concentration using Lenti-X Concentrator was performed with the manufacturer’s protocol and dissolved in Opti-MEM (Gibco #31985062) to yield a 10- or 15-fold concentration. The concentrated virus samples were stored at –80°C.

##### Transduction of HEK293T and HeLa cells

Cells were seeded on 6-well cell culture plate wells at a density of ∼2 x 10^5^ cells/well with 2 mL of the culture medium one day before transduction. A total of 1,000 µL transduction mix containing 1 µL of 2 µg/mL Polybrene (Sigma-Aldrich #TR-1003), recombinant lentivirus, and the cell culture medium was applied to each well alongside non-virus controls. One day after transduction, cells were trypsinized and plated on 96-well cell culture plate wells at a density of ∼5 x 10^3^ cells/well for a virus titer measurement. The next day, the culture medium was exchanged with a fresh medium containing 2.0 µg/mL Puromycin (Gibco #A1113803) or 5.0 µg/mL Blasticidin S (Wako #029-18701) to select the infected cells for two to five days. After drug selection, cell viability was measured using CellTiter-Glo (Promega #G7570) according to the manufacturer’s protocol. The luminescence was quantified using Infinite 200 PRO plate reader (TECAN). The background luminescence from wells without cell samples was used to subtract the signals. For each condition, the multiplicity of infection (MOI) was determined by the fraction of the survived cells compared to the non-selective condition control. The samples with an MOI close to but not over 0.1 was used for the following analyses, where most of the selected cells were expected to have a single viral integration according to Poisson statistics.

##### Transduction of mESCs

Cells were seeded on 6-well cell culture plate wells at a density of ∼2 x 10^5^ cells/well with 2 mL of the culture medium one day before transduction. For cell transduction, recombinant virus samples with volume of 10–100 µL were thawed on ice, mixed with 1.5 µL of 8 µg/mL Polybrene (Sigma-Aldrich #TR-1003) and 1.5 mL of fresh cell culture medium, and applied to the cells. To select the transduced cells, the culture media was exchanged with a fresh media containing 1.0 µg/mL Puromycin (Gibco #A1113803) two days after infection, followed by incubation for three days. Survived cells were detached, and cell counts were measured using Automated Cell Counter TC20 (BioRad). The MOI was determined by the fraction of the survived cells compared to the non-selective condition control. The samples with an MOI close to but not over 0.1 was used for the following analyses.

##### Transduction of hPSCs

Cells were seeded on 6-well cell culture plate wells at a density of ∼1 x 10^5^ cells/well with 2 mL of the culture medium one day before transduction. For cell transduction, recombinant virus samples with volume of 10–100 µL were thawed on ice, mixed with 1.5 µL of 8 µg/mL Polybrene (Sigma-Aldrich #TR-1003) and 1.5 mL of fresh cell culture medium, and applied to the cells. After 48 hours of infection, the culture media was exchanged with a fresh media containing 1.0 µg/mL of Puromycin (Gibco #A1113803) for three days. The reporter-integrated cells were dissociated into single cells and subjected to flow cytometry cell sorting to enrich EGFP-negative cells. The sorted cells were maintained with StemFit AK02N culture media (REPROCELL #RCAK02N).

#### Preparing cells with stably integrated Target-AID

##### mESCs

The mESC line with stably integrated Target-AID was established by electroporation using NEPA21 Super Electroporator (NEPAGENE). After detaching cells from cell culture plate wells, ∼2 x 10^6^ cells were mixed with 100 µL of Opti-MEM (Gibco #31985062), 2.0 µg of pNM1325, and 0.7 µg of a Super piggyBac transposase vector (SBI #PB210PA-1), and transferred to an electroporation cuvette (NEPAGENE #EC-002S). The electroporation was done by two poring pulses of positive electric polarity with 115 V for 5 ms with 50-ms intervals and 10% decay rate, and five transfer pulses each for positive and negative electric polarities with 20 V for 50 ms with 50-ms intervals and 40% the decay rate. After electroporation, cells were transferred to a 10-cm cell culture dish with a fresh culture medium. The medium was exchanged with a fresh medium one day after electroporation. Two days post electroporation, the medium was exchanged again with medium containing 5 µg/mL of Blasticidin S (Wako #029-18701) to select cells with stable integration. The cells were incubated for about two weeks in the selection medium.

##### hPSCs

To establish an hPSC line with stably integrated Target-AID, CA1 cells were seeded on 24-well cell culture plate wells at a density of ∼5 x 10^4^ cells/well with 1 mL of the culture medium one day before transfection. The next day, the medium was exchanged to remove Y-27632. The transfection mix was prepared by combining 450 ng of pNM1325 (CAGp-Target-AID-2A-Blast), 50 ng of a hyperactive PiggyBac transposase plasmid, 1 µL of Lipofectamine Stem Transfection Reagent (Invitrogen #STEM00001), and 49 µL of Opti-MEM (Gibco #31985062) and applied to the wells after 10 min incubation. The next day, the culture medium was exchanged with a fresh medium to remove the transfection reagent. Three days after transfection, the medium was exchanged with a fresh medium containing 5 µg/mL of Blasticidin S to start selection for 24 hours. Another selection was performed two days until confluent and cells were passaged into a new cell culture plate. One extra selection was conducted to ensure a positive selection of the cells.

#### Transient plasmid delivery

##### Transfection of HEK293T and HeLa cells

Cells were seeded on 24-well cell culture plate wells at a density of ∼5 x 10^4^ cells/well with 500 µL of the culture medium or 6-well cell culture plate wells at a density of ∼2 x 10^5^ cells/well with 2,000 µL of the culture medium one day before transfection. For regular transfection in a 24-well plate, a total of 400 ng of plasmids (3:1 volume of Cas9-based enzyme plasmid to gRNA plasmid when they were mixed), 1.2 µL of 1 mg/mL PEI MAX (Polyscience #24765), and 100 µL of 1x PBS were mixed, incubated for 5 or 10 min at room temperature, and applied to each well. The dose-dependent activation assay with the different Target-AID expression plasmid was performed using a 24-well plate with the plasmid amount per well ranging from 50–800 ng and 1 mg/mL of PEI MAX whose volume was adjusted to be 3 µL per 1 µg of the plasmid. For isolating barcoded HEK293T cells using CloneSelect C→T, we used 6-well cell culture plates, and a total of 800 ng of plasmids encoding both Target-AID and gRNA were combined with 2.5 µL of 1 mg/mL PEI MAX (Polyscience #24765) and 200 µL of 1x PBS and applied to each well after 5–10 min incubation at room temperature.

##### Transfection of mESCs

Cells were seeded on 48-well cell culture plate wells at a density of ∼6 x 10^4^ cells with 200 µL of the culture medium. For each reaction, a total of 200 ng of plasmids were diluted in 20 µL of Opti-MEM (Gibco #31985062), and 0.6 µL of Lipofectamine 2000 (Invitrogen #11668019) was combined with 19.4 µL of Opti-MEM (Gibco #31985062) as a transfection mix. The plasmid and transfection mix were then combined and applied to each well after 5 min incubation at room temperature.

##### Electroporation of hPSCs

For the gRNA-dependent reporter activation of the barcoded CA1 hPSCs with the stably integrated Target-AID, we used Neon Transfection System (Invitrogen MPK5000) to deliver the gRNA plasmid by electroporation. Cells were detached from cell culture plate wells, and ∼1 x 10^5^ cells were mixed with 100 µL of Neon Resuspension Buffer and 2.0 µg of gRNA plasmid. The electroporation was done by 1,200 V for 30 ms with one pulse. In the co-delivery of Target-AID and gRNA expression plasmids to the barcoded H1 hPSCs, ∼1 x 10^5^ cells were mixed with 100 µL of Neon Resuspension Buffer, 3.0 µg of Target-AID plasmid, and 3.0 µg of gRNA plasmid, and the electroporation was done by 1,200 V for 20 ms with two pulses.

#### Preparing microscope imaging samples

For imaging HEK293T cells, 25 µL of 0.1 mg/mL Hoechst 33342 (Invitrogen #H3570) dissolved in DMEM was directly added to each well of 24-well cell culture plates three days after transfection for nuclear counterstaining. The specimens were incubated at room temperature for 10 min, followed by removal of the culture medium. Cells were gently washed with fresh 500-µL of DMEM once and filled with 500 µL of fresh DMEM before imaging. For imaging HeLa cells, cells were first washed with 500 µL of 1x PBS, added 25 µL of 0.1 mg/mL Hoechst 33342 dissolved in 475 µL of DMEM and incubated at room temperature for 10 min. Cells were gently washed with 500 µL of 1x PBS and filled with 500 µL of fresh DMEM. For mESCs, 5.0 µg/mL Hoechst 33342 (Invitrogen #H3570) dissolved in the cell culture medium was directly added to each well and incubated at room temperature for 10 min to proceed with imaging. All live cell imaging was performed using BZ-X710 (Keyence), InCellAnalyzer 6000 (GE Healthcare), or IX83 (Olympus) with a 4x, 10x, or 20x objective lens. The contrast and brightness of the images obtained in a single batch of the experiment were uniformly adjusted using ImageMagick (Version 7.1.0-20) or Fiji (Version 1.0).

#### Flow cytometry analysis

Cells were detached by 0.25 w/v% Trypsin-EDTA (Wako #201-18841), incubated at 37°C for 5 min, collected into a 1.5-mL tube or a 96-well round-bottom plate, and centrifuged at 1,000 rpm and room temperature for 5 min. After aspirating the supernatant, cell pellets were gently resuspended with 150– 500 µL of ice-cold FACS buffer consisting of 2% FBS in 1x PBS. The samples were immediately placed on ice until flow cytometry analysis. The flow cytometry analysis was performed with BD FACSVerse Cell Analyzer (BD Biosciences) or CytoFLEX Flow Cytometer (Beckman Coulter). The samples were mixed gently by pipetting or vortexing immediately before the analysis. Approximately 10,000–20,000 raw events were acquired for each sample. The data analysis was performed with custom R scripts using flowWorkspace (version 0.5.40) (https://github.com/RGLab/flowWorkspace), flowCore (version 1.11.20) (https://github.com/RGLab/flowCore) and CytoExploreR (version 1.1.00) (https://github.com/DillonHammill/CytoExploreR) or a Python package FlowCytometryTools (version 0.5.0) (https://github.com/eyurtsev/FlowCytometryTools). Codes are available at https://github.com/yachielab/CloneSelect_v1/tree/main/FACS.

#### Flow cytometry cell sorting

##### HEK293T cells

Four days after the transfection of Target-AID and gRNA plasmids for barcode-specific cell isolation, cells were detached by 0.25 w/v% Trypsin-EDTA (Wako #201-18841), incubated at 37°C for 5 min, collected into a 1.5-mL tube, and centrifuged at 1,000 rpm and room temperature for 5 min, followed by resuspension into a 5-mL polystyrene round-bottom tube (FALCON) containing 150–500 µL of 1% FBS in 1x PBS. The cell suspension was immediately placed on ice until sorting. The sorting was performed using BD FACSJazz (BD Biosciences) with 1.0 Drop Single Sort mode. Cells were first gated using FSC-A and SSC-A, and the gate for EGFP+ cells was determined to obtain those having high FITC-A intensities that were not observed in a control cell sample transfected with Target-AID and NT gRNA plasmids. EGFP+ cells were sorted into 8-strip PCR tubes (Nippon Genetics #FG-018WF), each containing 2.5 µL of 1x PBS. For good cell recovery, the cell destination position in the collecting tube was adjusted manually for each sample. The sample was immediately placed on an ice-cold 96-well aluminum block. Although the EGFP+ cell rate varied across the samples, approximately 50–600 EGFP+ cells were recovered from each experiment.

##### mESCs

Three days after the transduction of a query gRNA, each cell sample was expanded in a 10-cm cell culture dish. Cells were detached by 0.25 w/v% Trypsin-EDTA (Gibco #25200072), incubated at 37°C for 5 min, collected into a 1.5-mL tube, and centrifuged at 1,000 rpm and room temperature for 5 min, followed by resuspension to ∼1 x 10^6^ cells in 1x PBS containing 2% FBS in a 5-mL polystyrene round-bottom tube (Falcon #352054). The cell suspension was immediately placed on ice until sorting. The sorting was performed using MoFlo Astrios EQ Cell Sorter (Beckman Coulter). Cells were first gated using FSC-A and SSC-A, and the gate for EGFP+ cells was determined to obtain those having high FITC-A intensities that were not observed in a non-transduction control cell sample. EGFP+ positive cells were single-cell sorted into 96-well plate wells, with the rest sorted in bulk to a single well of 96-well plate, each with 100 µL of the mESC culture media. Approximately 100–1,000 EGFP+ cells were recovered from each experiment, except for Clone 153, for which EGFP+ cells above the gating threshold could not be observed.

#### Barcode sequencing library preparation

##### CloneSelect C→T plasmid library and barcoded cell population

To identify barcodes of the CloneSelect C→T plasmid library by high-throughput sequencing, ∼10 ng of plasmid DNA (∼1.0 x 109 molecules) was used as a PCR template. To identify barcodes of the initial barcoded HEK293Ta cell population, genomic DNA was purified using NucleoSpin Tissue (Macherey Nagel #740952) according to the manufacturer’s protocol, and 119 ng of the extracted genomic DNA (4 x 10^4^ molecules; 400-fold to the estimated barcode complexity) was used as a PCR template. The sequencing libraries were prepared by a two-step PCR method. The first-round PCR was performed in triplicate, each in 20 µL volume, composed of template DNA, 0.5 µL each of 20 µM forward (SI#682) and reverse (SI#683) primers, 0.2 µL of Phusion High-Fidelity DNA Polymerase (NEB #M0530), 4 µL of Phusion HF Buffer (NEB #B0518S), 2 µL of 2 mM dNTPs (Takara #4025), and 0.6 µL of 100% DMSO (NEB #12611P), with the following thermal cycle condition: 98°C for 10 s, 30 cycles of 98°C for 10 s, 61°C for 10 s, and 72°C for 30 s, and then 72°C for 5 min for the final extension. Each PCR product was size-selected using 2% agarose gel, purified using PCR/Gel Extraction Kit (Nippon Genetics #FG-91302). To provide Illumina sequencing adapters and custom indices, the second-round PCR was performed to amplify each first-round PCR replicate product in 20 µL volume, composed of 2.5 ng of the 1st PCR product, 1 µL each of 10 µM P5 and P7 custom index primers, 0.2 µL of Phusion High-Fidelity DNA Polymerase (NEB #M0530), 4 µL of Phusion HF Buffer (NEB #B0518S), 2 µL of 2 mM dNTPs (Takara #4025), and 0.6 µL of 100% DMSO (NEB #12611P), with the following thermal cycle condition: 98°C for 10 s, 20 cycles of 98°C for 10 s, 61°C for 10 s, and 72°C for 30 s, and then 72°C for 5 min for the final extension. Custom indices assigned to the second-round PCR products can be found in Table S3. The second-round PCR products were size-selected and purified using PCR/Gel Extraction Kit (Nippon Genetics #FG-91302). The samples were pooled, quantified by qPCR using KAPA Library Quantification Kit Illumina (KAPA BIOSYSTEMS #KK4824), and analyzed by paired-end sequencing using Illumina MiSeq.

##### Sorted HEK293T cells

For the amplicon sequencing-based identification of barcodes in low-volume cells sorted after gRNA-dependent barcode-specific clone isolation using CloneSelect C→T, we prepared a cell lysate for each sample as a PCR template. The sequencing library of each sample was prepared by modifying the two-step PCR protocol described for purified DNA as a PCR template. Cells in 8-strip PCR tubes were first incubated with 2.0 µL of lysis buffer, including 600 mM KOH, 10 mM EDTA, and 100 mM DTT. The samples were then neutralized with 2.0 µL of neutralization buffer composed of 0.4 µL of 1M Tris-HCl and 1.6 µL of 3 M HCl. The first-round PCR was performed by replacing the template with 2.0 µL of the cell lysate. Although we did not observe visible gel electrophoresis bands for the first-round PCR products, the PCR product of the expected size were size-selected using 2% agarose gel, purified and eluted into 15 µL of ddH2O using PCR/Gel Extraction Kit (Nippon Genetics #FG-91302). The PCR products were quantified using Quant-iT PicoGreen dsDNA Assay Kit (Thermo Fisher Scientific #P7589) and Infinite 200 PRO plate reader (TECAN) using Tecan i-control software (version 1.10.4.0). The second-round PCR was performed for 2.0 ng of the first-round PCR product. Custom indices assigned to the second-round PCR products can be found in Table S3. The second-round PCR products were size-selected and purified using PCR/Gel Extraction Kit (Nippon Genetics #FG-91302). The samples were pooled into a DNA LoBind 1.5-mL tube (Eppendorf #13-698-791), quantified by qPCR using KAPA Library Quantification Kit Illumina (KAPA BIOSYSTEMS #KK4824), and analyzed by paired-end sequencing using Illumina MiSeq.

##### Uptag-dntag combination reference database

To identify the uptag-dntag combination reference database of the mESC population tagged using the scCloneSelect barcode library, genomic DNA was first extracted from 1 x 10^5^ cells using NucleoSpin Tissue (MACHEREY-NAGEL #740952) according to the manufacturer’s protocol. The sequencing libraries were prepared by a two-step PCR method. A total of 50 ng genomic DNA was used for each PCR reaction. The first-round PCR was performed in duplicate, each in 20 µL volume, composed of template DNA, 0.7 µL each of 10 µM forward (SI#682) and reverse (RS#250) primers, 0.2 µL of Phusion High-Fidelity DNA Polymerase (NEB #M0530), 4.5 µL of Phusion HF Buffer (NEB #B0518S), 1.6 µL of 2.5 mM dNTPs (NEB #N0447), with the following thermal cycle condition: 98°C for 10 s, 30 cycles of 98°C for 10 s, 60°C for 10 s, and 72°C for 2 min, and then 72°C for 5 min for the final extension. Each PCR product was size-selected using 2% agarose gel, purified, and eluted into 20 µL of ddH2O using PCR/Gel Extraction Kit (Nippon Genetics #FG-91302). To provide Illumina sequencing adapters and custom indices, the second-round PCR was performed to amplify each first-round PCR replicate product in 20 µL volume, composed of 20-fold dilution of the first-round PCR product, 0.7 µL each of 10 µM P5 and P7 custom index primers, 0.2 µL of Phusion High-Fidelity DNA Polymerase (NEB #M0530), 4.5 µL of Phusion HF Buffer (NEB #B0518S), 1.6 µL of 2.5 mM dNTPs (NEB #N0447), with the following thermal cycle condition: 98°C for 10 s, 20 cycles of 98°C for 10 s, 60°C for 10 s, and 72°C for 30 s, and then 72°C for 5 min for the final extension. Custom indices assigned to the second-round PCR products can be found in Table S3. The second-round PCR products were size-selected and purified using PCR/Gel Extraction Kit (Nippon Genetics #FG-91302). The samples were pooled, quantified by qPCR using KAPA Library Quantification Kit Illumina (KAPA BIOSYSTEMS #KK4824), and analyzed by paired-end sequencing using Illumina MiSeq.

##### Sorted mESCs

For the amplicon sequencing-based identification of barcodes in cells sorted after gRNA-dependent barcode-specific clone isolation using scCloneSelect, we prepared a cell lysate for each sample as a PCR template. The sequencing library of each sample was prepared by modifying the two-step PCR method described for the identification of the uptag-dntag combination reference database. Sorted mESC samples were expanded in 96-well culture plate wells until confluent. After aspirating the culture medium, 20 µL of 50 mM NaOH was added to each well and transferred into a 96-well PCR plate for direct cell lysis. The samples were then heated at 95°C for 15 min, followed by cooling down on ice. The samples were neutralized with 2.0 µL of 1 M Tris-HCl (pH 8.0). The first-round PCR was performed in 40 µL volume replacing the template with 3.5 µL of cell lysate. The second-round PCR was performed in 20 µL volume replacing the template with a 10-fold dilution of the first-round PCR product. Custom indices assigned to the second-round PCR products can be found in Table S3. The second-round PCR products were size-selected and purified using GeneJET Gel Extraction Kit (Thermo Fisher Scientific #K0691). The samples were pooled into a DNA LoBind 1.5-mL tube (Eppendorf #0030108051), quantified by qPCR using KAPA Library Quantification Kit Illumina (KAPA BIOSYSTEMS #KK4824), and analyzed by paired-end sequencing using Illumina HiSeq2500.

##### Reamplification of scCloneSelect dntags from Drop-seq library

To increase the sensitivity of identifying dntags associated with single cells and their transcriptome profiles, we selectively reamplified the DNA fragments encoding dntags and cell IDs from the intermediate Tn5 transposon-fragmented library of the Drop-seq process and sequenced them separately. The reamplification PCR was performed in 20 µL volume, composed of 1 ng of the cDNA library quantified using TapeStation with High Sensitivity D5000 ScreenTape (Agilent #5067-559 and #5067-5593), 0.7 µL each of 20 µM forward primer P5-TSO_Hybrid (*45*) and reverse primer SI#682, 0.2 µL of Phusion High-Fidelity DNA Polymerase (NEB #M0530), 4.5 µL of 5x Phusion HF Buffer (NEB #B0518), and 1.6 µL of 2.5 mM dNTPs (NEB #N0447) with the following thermal cycle condition: 95°C for 30 s, 30 cycles of 98°C for 30 s, 60°C for 10 s, and 72°C for 2 min, and 72°C for 5 min for the final extension. The first-round PCR product was purified and eluted into 20 µL of ddH2O using GeneJET Gel Extraction Kit (Thermo Fisher Scientific #K0691). The second-round PCR was then reamplified in 20 µL volume, composed of 10-fold dilution of the first-round PCR product, 0.7 µL each of 10 µM P5 and P7 custom dual index primers, 0.2 µL of Phusion High-Fidelity DNA Polymerase (NEB #M0530), 4.5 µL of 5x Phusion HF Buffer (NEB #B0518), 1.6 µL of 2.5 mM dNTPs (NEB #N0447), with the following thermal cycle condition: 95°C for 30 s, 15 cycles of 98°C for 10 s, 65°C for 10 s, and 72°C for 2 min, and 72°C for 5 min for the final extension. Custom indices assigned to the second-round PCR products can be found in Table S3. The second-round PCR products were size-selected using 2% agarose gel and purified using GeneJET Gel Extraction Kit (Thermo Fisher Scientific #K0691). The samples were pooled, quantified by qPCR using KAPA Library Quantification Kit Illumina (KAPA BIOSYSTEMS #KK4824), and analyzed by paired-end sequencing using Illumina HiSeq2500.

#### Drop-seq

scRNA-seq was performed by Drop-seq. The Drop-seq platform was set up using devices manufactured by Dolomite Bio following the manufacturer’s protocol. The microfluidic devices were fabricated by YODAKA Co., Ltd. Cell samples were prepared at a concentration of ∼2 x 10^5^ cells/mL for analysis. Sequencing libraries were prepared following the original Drop-seq paper (*45*). In brief, following emulsion breakage and reverse transcription, “single-cell transcriptomes attached to microparticles” (STAMPs) were washed and proceeded to the Exonuclease I (NEB #M0293L) treatment. A total of ∼2,000 STAMPs were used for the whole cDNA amplification of each sample. After second-strand synthesis, the library DNA was purified using AMPure XP beads (Beckman Coulter #A63881), quantified using TapeStation with High Sensitivity D5000 ScreenTape (Agilent #5067-5592 and #5067-5593), and fragmented by Tn5 transposon using Nextera XT DNA Library Preparation Kit (Illumina #FC-131-1024) according to the manufacturer’s protocol. The tagmented sequencing library was then purified using AMPure XP beads (Beckman Coulter #A63881) and quantified by TapeStation with High Sensitivity D5000 ScreenTape (Agilent #5067-5592 and #5067-5593). We confirmed each average library size to be ∼500 bp. Multiple scRNA-seq libraries were pooled and subjected to high-throughput sequencing using Illumina MiSeq or HiSeq2500. The sequencing library index information can be found in Table S3.

#### mESC differentiation assay

For the establishment of a barcoded mESC population, cells were seeded on 6-well cell culture plate wells at a density of ∼2 x 10^5^ cells/well with 2 mL of the culture medium containing 10^3^ units/mL ESGRO Recombinant Mouse Leukemia inhibitory factor (LIF) (Millipore #ESG1107) and 2i (1.0 µM PD0325901 Toris #4423 and 3.0 µM CHIR99021 Wako #038-23101). The next day, cells were transduced with 500 µL of 15-fold concentrated barcoded lentivirus pool of scCloneSelect. Two days after, the culture medium was exchanged with a fresh medium containing 1.0 µg/mL of Puromycin (Gibco #A1113803). To restrict the clone complexity for the downstream cell differentiation and clone isolation assays, a clonal population bottleneck was created where ∼1,000 cells were seeded on a 6-well cell culture plate well and cultured for ten days. Cells were then split as follows: ∼1 x 10^4^ cells were seeded into the culture medium with LIF and 2i (LIF+2i+), ∼1 x 10^4^ cells were seeded into the culture medium without LIF or 2i (LIF–2i–), two samples of ∼1 x 10^5^ cells each were used for the identification of uptag-dntag combination reference database, and five replicates of ∼1 x 10^5^ cells were stored at –80°C using CELLBANKER 1 freeze media (ZENOAQ #11910). Four days later, the cell cultures of the LIF+2i+ and LIF–2i– conditions were subjected to scRNA-seq.

#### RT-PCR-based analysis of barcode transcription

The transcription of polyadenylated scCloneSelect barcode products was analyzed by RT-PCR and gel electrophoresis. Total RNA was first extracted using ISOSPIN Cell & Tissue RNA Kit (Nippon Gene #314-08211) according to the manufacturer’s protocol. The RNA sample was then subjected to DNase I treatment (Takara #2270B) to remove DNA and purified again using ISOSPIN Cell & Tissue RNA Kit (Nippon Gene #314-08211). The first-strand cDNA was synthesized using High-Capacity cDNA Reverse Transcription Kit (Applied Biosciences #4368814). The reaction was performed in 10 µL volume, composed of 5 µL of DNase I treated RNA (∼1 µg), 0.5 µL of 100 µM oligo dT primer SI#4, 0.5 µL of MultiScribe Reverse Transcriptase, 1 µL of 10x RT buffer, 0.4 µL of 100 mM dNTP, and 0.5 µL of RNAse Inhibitor (Applied Biosciences #N8080119), with the following thermal cycler condition: 25°C for 10 min, 37°C for 12 min, and 85°C for 5 min. Lastly, the transcription of the target barcode was tested by PCR along with a GAPDH control. The PCR was performed in 20 µL volume, composed of 2 µL of 50-fold diluted first-strand cDNA, 2.8 µL total of a primer pair SI#116–SI#7 to amplify the dntag or a primer pair RS#507–RS#508 to amplify GAPDH, 0.2 µL of Phusion DNA Polymerase (NEB #M0530S), 4 µL of 5x Phusion HF Buffer (NEB #B0518), and 1.6 µL of 2.5 mM dNTPs (NEB #N0447), with the following thermal cycler condition: 98°C for 30 s, 30 cycles of 98°C for 10 s, 60°C for 10 s, and 72°C for 30 s, and then 72°C for 5 min for the final extension. The resulting PCR products were analyzed with 2% agarose gel.

### Yeast CloneSelect

#### Strains

*Saccharomyces cerevisiae* BY4741 (*MATa his3Δ0 leu2Δ0 met15Δ0 ura3Δ0*) was used for the experiments using yeast CloneSelect system.

#### Yeast CloneSelect library

To generate the yeast CloneSelect barcode library, a semi-random oligo pool KI#200 encoding 5′-CCGWSNSWSNSWSNSWSNSNGTG-3′ was first chemically synthesized (Table S2) and amplified by PCR in 40 µL volume, composed of 2 µL of 0.01 µM template, 2 µL each of 10 µM forward primer SI#368 and 10 µM reverse primer SI#369, 0.8 µL of Phusion High-fidelity DNA Polymerase (NEB #M0530S), 8 µL of 5x Phusion HF Buffer (NEB #B0518S), and 2 µL of 2 mM dNTPs, with the following thermal cycle condition: 98°C for 30 s, 35 cycles of 98°C for 10 s, 68°C for 20 s, and 72°C for 5 s, and then 72°C for 5 min for the final extension. The PCR product was analyzed with 2% agarose gel and size-selected and purified using FastGene PCR/Gel Extraction Kit (Nippon Genetics #FG-91302). The purified barcode fragment was then assembled into the cloning backbone plasmid pKI110 by Golden Gate Assembly (*66*) using BsmBI (NEB #R0580S). We performed two assembly reactions, each in 25 µL volume, composed of 500 fmol barcode fragments, 50 fmol backbone plasmid, 0.5 µL of BsmBI (NEB #R0580S), 0.5 µL of T4 DNA Ligase (Nippon Gene #317-00406), 2.5 µL of 10x T4 DNA Ligation Reaction Buffer (NEB #B0202S), and 0.125 µL of 60 mg/mL BSA (NEB #B9001S) with the following thermal cycle condition: 15 cycles of 37°C for 5 min and 20°C for 5 min, and then 55°C for 30 min for complete backbone digestion. For the bacterial transformation, 5 µL of the assembly product was used to transform 50 µL of DH5α chemically competent cells (NEB #C2987I) according to the manufacturer’s high-efficiency transformation protocol. Following one-hour outgrowth in 1 mL of SOC medium (NEB #B9020S) at 37°C, cells were plated on a total of four LB agar plates containing 100 µg/mL Ampicillin (Wako #014-23302). The cell sample was also diluted and spread on the selective plates to estimate the clone complexity. To check assembly efficiency, random clones were isolated to perform genotyping PCR with a primer pair of KI#169 and KI#170, validating each clone to have an expected barcode insert. To construct Pool-100, 100 colonies were isolated, dissolved in 80 µL each of LB medium containing 100 µg/mL Ampicillin (Wako #014-23302), combined 5 µL each, and cultured overnight at 37°C. The plasmid DNA was extracted using FastGene Plasmid Mini Kit (Nippon Genetics #FG-90502). Pool-1580 was constructed by scraping and harvesting colonies from a plate with colony forming units close to 1,000 by adding 1.5 mL of LB media containing 100 µg/mL Ampicillin (Wako #014-23302). The resulting cell samples were centrifuged at 13,000 rpm for 2 min to discard supernatant. The plasmid DNA pools were then purified from the collected cells using FastGene PCR/Gel Extraction Kit (Nippon Genetics #FG-91302).

#### Barcoding of cells and introduction of genome editing reagents

For the barcoding of cells and introduction of genome editing reagents by transformation, we used the Frozen-EZ Yeast Transformation II kit (Zymo Research #T2001) with slight modifications. Cells were first pre-cultured in 5 mL of YPDA or SC-Dropout medium (complying with the auxotrophic requirement for plasmid maintenance in the host cells) in a cell culture tube rotating overnight at 30°C. The next day, cells were cultured in 5 mL of a fresh YPDA medium with a starting OD_600 nm_ of 0.3 and incubated until the OD_600 nm_ reached 0.8–1.0. After making competent cells according to the manufacturer’s protocol, plasmid DNA and 50 µL of competent cells were added into a 1.5-mL tube, mixed thoroughly with 500 µL of EZ3 solution according to the manufacturer’s protocol, and incubated at 30°C for an hour with rotation. The cell sample was centrifuged at 13,000 rpm for 2 min, and the supernatant was discarded, followed by the addition of 2.5 mL of YPDA medium for recovery. After two-hour outgrowth at 30°C with rotation, cells were centrifuged to remove the medium and washed with 1,000 µL of 1x TE twice. Cells were then spread on SC–Dropout agar plates and incubated for two–four days at 30°C.

##### Barcoding of cells

When the background BY4741 cells were transformed using the barcode plasmid library containing *HIS3* marker, YPDA medium was used for pre-culturing and SC–His+Ade plates for selecting transformants. For the pooled barcoding of cells, the reaction was scaled to transform 250 µL of competent cells using 200 ng of plasmid in total, and colonies formed on the selective plates were pooled and collected by scraping with 3–4 mL of SC–His+Ade. For the barcoding of cells using a single barcode plasmid clone, 200 ng of plasmid was used to transform 15 µL of competent cells.

##### Introduction of genome editing reagents

When cells harboring the barcode plasmid containing *HIS3* marker were subjected to a clone isolation, cells were transformed twice, first with the constitutively active Target-AID plasmid pKI086 containing *LEU2* marker and next with the targeting gRNA expression plasmid encoding *URA3* marker. For the first transformation, we used SC–His+Ade medium for pre-culturing and SC–His–Leu+Ade plates for selecting transformants. For the second transformation, SC– His–Leu+Ade medium for pre-culturing and SC–His–Leu–Ura+Ade for selecting transformants. When the background BY4741 cells were transformed using one of the galactose-inducible Cas9-based enzyme plasmids (Cas9, dCas9, dCas9-PmCDA1, dCas9-PmCDA1-UGI, nCas9, nCas9-PmCDA1, and nCas9-PmCDA1-UGI) containing *LEU2* marker and a CAN1-targeting gRNA plasmid containing *URA3* marker, we used YPDA medium for pre-culturing and SC–Leu–Ura+Ade plates for selecting transformants. For the barcode-specific reporter activation in a complex barcoded population, the reaction was scaled to transform 250 µL of competent cells using 200 ng of the enzyme plasmid and 200 ng of the targeting gRNA plasmid in total. For the other small-scale transformations, 200 ng of the effector plasmid and 200 ng of the targeting gRNA plasmid were used to transform 15 µL of competent cells.

#### Barcode sequencing library preparation

To identify barcodes of the yeast CloneSelect plasmids introduced to the cells by high-throughput sequencing, we first extracted and purified the barcode plasmids. We first centrifuged cells at 15,000 rpm for 3 min, followed by discarding the supernatant. The cell pellet was resuspended with 20 µL of Zymolyase Buffer, containing 2.5 mg/mL Zymolyase (Zymo Research #E1005), 10 mM Sodium phosphate, and 1.2 M Sorbitol, and 500 µL of Solution I Buffer (Supplied in Zymolyase #E1005), containing 0.1 M EDTA and 1 M Sorbitol. The sample was incubated at 37°C for one hour and centrifuged at 15,000 rpm for 1 min, followed by discarding the supernatant. The cell lysate sample was then incubated with 250 µL of Solution II Buffer (Supplied in Zymolyase #E1005), containing 20 mM EDTA and 50 mM Tris-HCl, and 1% SDS, at 65°C for 30 min, followed by the addition of 100 µL of 5 M potassium acetate. The sample was further incubated on ice for 30 min and centrifuged at 15,000 rpm for 3 min. The supernatant was transferred into a 1.5-mL tube, and the plasmid DNA was precipitated with the addition of 400 µL of isopropanol, followed by cleanup with 400 µL of 70% ethanol. The resulting DNA pellet was resuspended with 50 µL of ddH2O containing 10 µg/mL RNase and incubated at 65°C for 10 min. Next, the barcode sequencing libraries were prepared by a two-step PCR method. The first-round PCR was performed in triplicate, each in 40 µL volume, composed of 1.0 µg of template DNA, 1 µL each of 10 µM forward primer KI#169 and 10 µM reverse primer KI#289, 0.4 µL of Phusion High-Fidelity DNA Polymerase (NEB #M0530S), 8 µL of Phusion HF Buffer (NEB #B0518S), and 0.8 µL of 10 mM dNTPs (Takara #4030), with the following thermal cycle condition: 98°C for 30 s, 20 cycles of 98°C for 10 s, 61°C for 20 s, and 72°C for 25 s, and then 72°C for 5 min for the final extension. Each PCR product was size-selected using 2% agarose gel, purified, and eluted into 50 µL of ddH2O using PCR/Gel Extraction Kit (Nippon Genetics #FG-91302). To provide Illumina sequencing adapters and custom indices, the second-round PCR was performed to amplify each first-round PCR replicate product in 40 µL volume, composed of 2 µL of the first PCR product, 1 µL each of 10 µM P5 and P7 custom index primers, 0.4 µL of Phusion High-Fidelity DNA Polymerase (NEB #M0530S), 8 µL of 5x Phusion HF Buffer (NEB #B0518S), and 0.8 µL of 10 mM dNTPs (Takara #4030), with the following thermal cycle condition: 98°C for 30 s, 15 cycles of 98°C for 10 s and 60°C for 10 s, and 72°C for 1 min, and then 72°C for 5 min for the final extension. Custom indices assigned to the second-round PCR products can be found in Table S3. The second-round PCR products were size-selected and purified using PCR/Gel Extraction Kit (Nippon Genetics #FG-91302). The sequencing libraries were pooled, quantified by qPCR using KAPA Library Quantification Kit Illumina (KAPA BIOSYSTEMS #KK4824), pooled with equimolar ratios, and analyzed by paired-end sequencing using Illumina HiSeq2500.

#### Analysis of reporter activation efficiency

To analyze the efficiency of the gRNA-dependent barcode-specific mCherry reporter activation, we treated three independent barcoded cell samples, each by all three corresponding gRNAs (3x3 assay). Each sample was spread on SC–His–Leu–Ura+Ade agar plates, scraped and inoculated to a 1.5-mL tube containing 500 µL of SC–His–Leu–Ura+Ade medium, and cultured for two–four days at 30°C. The 20 µL of pre-cultured cell sample was mixed with 180 µL of SC–His–Leu–Ura+Ade media and transferred into a flat-bottom transparent 96-well plate (Greiner Bio-One #655090). Samples were then analyzed using Infinite 200 PRO plate reader (TECAN) using Tecan i-control software (version 1.10.4.0) to measure mCherry fluorescence intensities normalized by OD_595 nm_ values. For microscopic observation of cells, 2.5 µL of cell sample was transferred on a glass slide, gently covered with a glass coverslip, and observed using BZ-X710 (Keyence) with 20x and 40x objective lenses. We also directly measured the GTG→ATG conversion rate of each sample by high-throughput sequencing. Cells were scraped and harvested from the selective plates and lysed by DNAZol (COSMO BIO #DN127) according to the manufacturer’s protocol. The sequencing libraries were then prepared by a two-step PCR method. The first-round PCR was performed in 32 µL volume, composed of 1.6 µL of cell lysate, 1.6 µL each of 10 µM forward primer KI#168 and 10 µM reverse primer KI#169, 0.64 µL of Phusion High-Fidelity DNA Polymerase (NEB #M0530S), 6.4 µL of Phusion HF Buffer (NEB #B0518S), and 0.64 µL of 10 mM dNTPs (Takara #4030), with the following thermal cycle condition: 98°C for 30 s, 30 cycles of 98°C for 10 s, 61°C for 10 s, and 72°C for 1 min, and then 72°C for 5 min for the final extension. The remaining procedure was processed by the same protocols described for the barcode sequencing library preparation. Custom indices assigned to the second-round PCR products can be found in Table S3. The second-round PCR products were size-selected and purified using PCR/Gel Extraction Kit (Nippon Genetics #FG-91302). The samples were pooled, quantified by qPCR using KAPA Library Quantification Kit Illumina (KAPA BIOSYSTEMS #KK4824), and analyzed by paired-end sequencing using Illumina MiSeq.

#### Isolation and analysis of barcoded colonies

After the barcode-specific reporter activation of a complex population, cells from test and control conditions were spread on SC–His–Leu–Ura+Ade agar plates and imaged under a blue light illuminator FAS-IV (Nippon Genetics) to distinguish mCherry+ and mCherry– colonies. Colonies were isolated into 96-well cell culture plate wells containing 98 µL of SC–His–Leu–Ura+Ade and cultured overnight at 30°C. Samples were then analyzed using Infinite 200 PRO plate reader (TECAN) using Tecan i-control software (version 1.10.4.0) to measure mCherry fluorescence intensities normalized by OD_595 nm_ values. The same colony isolates were also subjected to Sanger sequencing to identify their barcode sequences and base editing outcomes. Using the same cell lysis, first-round PCR, and PCR cleanup protocols as in the analysis of the reporter activation efficiency, we obtained barcode DNA fragments from each sample. Each PCR product was analyzed by Sanger sequencing using a sequencing primer SI#658. The Sanger sequencing trace was analyzed using PySanger (https://github.com/ponnhide/PySanger).

#### Canavanine assay

Genome editing efficiencies of different Cas9-based genome editing enzymes (Cas9, dCas9, dCas9-PmCDA1, dCas9-PmCDA1-UGI, nCas9, nCas9-PmCDA1 and nCas9-PmCDA1-UGI) were estimated by Canavanine assay, where enzymes were introduced to the cells with a gRNA targeting an arginine transporter CAN1 and its knockout efficiency can be assessed by cell survival under the presence of a toxic arginine analog Canavanine. In this assay, Cas9-based enzyme expressions were regulated under a galactose-inducible GAL1/10 promoter. To induce genome editing, cells containing enzyme and gRNA plasmids were first cultured in SC–Leu–Ura medium containing 2% glucose at 30°C until saturation. Cells were then resuspended in SC–Leu–Ura medium containing 2% raffinose with 16-fold dilution and cultured at 30°C until saturation. Finally, cells were resuspended in SC–Leu–Ura medium containing 2% raffinose and 0.02% galactose with 32-fold dilution and cultured at 30°C for two days. Each sample was then spread on SC–Leu–Ura–Arg+Ade and SC–Leu–Ura–Arg+Ade containing 60 mg/ml canavanine plates and cultured at 30°C for two–four days to estimate colony forming units and perform spot assays. Furthermore, we scraped and harvested colonies from the SC–Leu–Ura–Arg+Ade control plates to extract genomic DNA samples for the mutation spectra assay by high-throughput sequencing. We first lysed 20 µL of cells at OD_600 nm_ of 1.0 using 100 µL of DNAzol (COSMO BIO #DN127) followed by incubation at room temperature for 15 min. The lysate was mixed thoroughly with 30 µL of 1M NaCl and 50 µL of 100% ethanol and centrifuged at 14,000 rpm for 10 min. The supernatant was discarded, and the pellet was washed with 550 µL of 70% ethanol. After air-drying, the sample was resuspended in 50 µL of ddH2O. We then prepared an amplicon sequencing library for each sample by a two-step PCR method in triplicate. The first-round PCR was in 40 µL volume, composed of 2 µL of template DNA, 2 µL each of 10 µM forward primer #KN85F3 and 10 µM reverse primer #KN85R2, 0.8 µL of Phusion High-Fidelity DNA Polymerase (NEB #M0530S), 8 µL of Phusion HF Buffer (NEB #B0518S), and 0.8 µL of 10 mM dNTPs (Takara #4030), with the following thermal cycle condition: 98°C for 30 s, 30 cycles of 98°C for 10 s, 60°C for 10 s, and 72°C for 1 min, and then 72°C for 5 min for the final extension. We also performed the same process using a primer pair #HO2F2–#HO2R2 to prepare control samples. The remaining procedure was processed by the same protocols described for the barcode sequencing library preparation. Custom indices assigned to the second-round PCR products can be found in Table S3. The second-round PCR products were size-selected and purified using PCR/Gel Extraction Kit (Nippon Genetics #FG-91302). The sequencing libraries were pooled, quantified by qPCR using KAPA Library Quantification Kit Illumina (KAPA BIOSYSTEMS #KK4824), pooled with equimolar ratios, and analyzed by paired-end sequencing using Illumina MiSeq.

### Bacterial CloneSelect

#### Preparation of cells for various bacterial CloneSelect systems

We prepared cell samples harboring single barcode plasmids for different variants of bacterial CloneSelect systems (Table S2). The plasmid for the EGFP reporter-based system was introduced to BL21(DE3) Competent *E. coli* cells (NEB #C2527I), and the plasmids for the Blasticidin and Zeocin resistance marker-based systems were introduced to T7 Express chemically competent cells (NEB #C2566I) according to the manufacturer’s high-efficiency transformation protocols. The transformants were selected on LB agar plates containing 100 µg/mL Ampicillin (Wako #014-23302) and/or 50 µg/mL Kanamycin (Wako #111-00344).

#### Bacterial CloneSelect library

To generate the bacterial CloneSelect barcode library for the Zeocin resistance marker, a semi-random oligo pool KI#405 encoding 5′-ATGCCGVNNVNNVNNVNNVNNTAA-3′ was chemically synthesized (Table S2), where a start codon ATG, the antisense strand sequence of 5′-CGG-3′ PAM and a quintuple repeat of VNN (V=non-T) were followed by a TAA stop codon. The VNN repeat restricts additional stop codons from appearing in-frame to the downstream reporter. The semi-random oligo pool was amplified by PCR in 20 µL volume, composed of 1 µL of 1 μM template, 1 µL each of 10 µM forward primer SI#368 and 10 µM reverse primer SI#369, 0.4 µL of Phusion High-fidelity DNA Polymerase (NEB #M0530L), 4 µL of 5x Phusion HF Buffer (NEB #B0518S), and 0.4 µL of 10 mM dNTPs, with the following thermal cycle condition: 98°C for 30 s, 20 cycles of 98°C for 10 s, 68°C for 20 s, and 72°C for 20 s, and then 72°C for 5 min for the final extension. The PCR product was analyzed with 2% agarose gel and size-selected and purified using FastGene PCR/Gel Extraction Kit (Nippon Genetics #FG-91302). The purified barcode fragment was then assembled into the cloning backbone plasmid pKI243 by Golden Gate Assembly using BsmBI (*66*). We performed the assembly in 12.5 µL volume, composed of 2.91 fmol of barcode fragments, 14.9 fmol of backbone plasmid, 0.25 µL of BsmBI (NEB #R0580L), 0.5 µL of T4 DNA Ligase (Nippon Gene #317-00406), 1.25 µL of 10x T4 DNA Ligation Reaction Buffer (NEB #B0202S), and 0.62 µL of 2 mg/mL BSA (NEB #B9001S) with the following thermal cycle condition: 15 cycles of 37°C for 5 min and 20°C for 5 min, and then 55°C for 30 min for complete backbone digestion. For the bacterial transformation, 3 µL of the assembly product was used to transform 65 µL of T7 Express chemically competent cells (NEB #C2566I) according to the manufacturer’s high-efficiency transformation protocol. Following one-hour outgrowth in 500 µL of SOC medium (NEB #B9020S) at 37°C, the cell sample was plated 250 µL each on three LB agar plates containing 100 µg/mL Ampicillin (Wako #014-23302). The cell sample was also diluted and plated on the selective plates to estimate the clone complexity. To check assembly quality and efficiency, 12 random clones were isolated and subjected to Sanger Sequencing, validating that 11 out of 12 clones showed an expected clone barcode insert. To construct Pool-100, 100 colonies were isolated, dissolved in 80 µL each of LB medium containing 100 µg/mL Ampicillin (Wako #014-23302), combined 5 µL each, and cultured overnight at 37°C. The plasmid DNA was extracted using FastGene Plasmid Mini Kit (Nippon Genetics #FG-90502). Pool-1550 was constructed by scraping and harvesting colonies from a plate with colony forming units close to 1,000 by adding 1.5 mL of LB media containing 100 µg/mL Ampicillin (Wako #014-23302). The barcode plasmid libraries were used to transform T7 Express chemically competent cells (NEB #C2566I) to establish barcoded *E. coli* cell populations.

#### Barcode sequencing library preparation

For each of the Pool-100 and Pool-1550 barcode plasmid libraries, the barcode sequencing library was prepared in triplicate by a two-step PCR method. The first-round PCR was performed in five separate PCR reactions, each in 40 µL volume, composed of 2.0 ng of plasmid template DNA, 1 µL each of 10 µM forward primer KI#403 and 10 µM reverse primer KI#404, 0.4 µL of Phusion High-Fidelity DNA Polymerase (NEB #M0530L), 8 µL of Phusion HF Buffer (NEB #B0518S), and 0.8 µL of 10 mM dNTPs (Takara #4030), with the following thermal cycle condition: 98°C for 30 s, 20 cycles of 98°C for 10 s, 54°C for 20 s, and 72°C for 25 s, and then 72°C for 5 min for the final extension. For each replicate, the five PCR reaction products were pooled, size-selected using 2% agarose gel, purified, and eluted into 30 µL of ddH2O using PCR/Gel Extraction Kit (Nippon Genetics #FG-91302). To provide Illumina sequencing adapters and custom indices, the second-round PCR was performed to amplify each first-round PCR replicate product in 40 µL volume, composed of 2 µL of the first PCR product, 1 µL each of 10 µM P5 and P7 custom index primers, 0.4 µL of Phusion High-Fidelity DNA Polymerase (NEB #M0530L), 8 µL of 5x Phusion HF Buffer (NEB #B0518S), and 0.8 µL of 10 mM dNTPs (Takara #4030), with the following thermal cycle condition: 98°C for 30 s, 15 cycles of 98°C for 10 s, 60°C for 10 s, and 72°C for 60 s, and then 72°C for 5 min for the final extension. Custom indices assigned to the second-round PCR products can be found in Table S3. The second-round PCR products were size-selected and purified using PCR/Gel Extraction Kit (Nippon Genetics #FG-91302). The sequencing libraries were pooled, quantified by qPCR using KAPA Library Quantification Kit Illumina (KAPA BIOSYSTEMS #KK4824), pooled with equimolar ratios, and analyzed by paired-end sequencing using Illumina MiSeq.

#### Introduction of genome editing reagents

To introduce a plasmid containing ABE-7.10 and gRNA with given induction systems to barcoded cells, we used Mix&Go! *E. coli* Transformation Kit (Zymo Research #T3001) according to the manufacturer’s protocol. To select successful transformants, the transformation reaction sample was plated on LB agar plates containing 100 µg/mL Ampicillin (Wako #014-23302) and 50 µg/mL Kanamycin (Wako #111-00344) and incubated overnight at 37°C. For the experiments that involved Arabinose (Sigma-Aldrich #A3256-10MG) and Isopropylthio-β-galactoside (IPTG) (ThermoFisher Scientific #15529019) inductions, cells were cultured in the medium containing 100 mM Arabinose and 0.1 mM IPTG overnight at 37°C prior to the analyses. For the barcoded cell isolation with the Zeocin-resistance marker-based system, we used a low-salt LB medium adjusted to pH 7.5 with 1 M NaOH (Nakalai #37421-05) for the Zeocin activity. We also decided to use 100 µg/mL Zeocin (Invitrogen #R25001) or 100 µg/mL Blasticidin S (Wako #029-18701) with no Arabinose or IPTG for genome editing and selecting reporter-activated cells, as leaky gene editing reagent expressions from the inducible promoters with the no inducer condition were sufficient and maximized the cell viability. The information on the genome editing plasmids used in this study can be found in Table S2.

#### Analysis of reporter activation efficiency

To analyze the efficiency of the gRNA-dependent barcode-specific EGFP-reporter activation, 200 µL of cell samples were transferred into a flat-bottom transparent 96-well plate (Greiner Bio-One #655090) and analyzed using Infinite 200 PRO plate reader (TECAN) using Tecan i-control software (version 1.10.4.0) to measure EGFP fluorescence intensities normalized by OD_595 nm_ values. For microscopic observation of cells, 2.5 µL of cell sample was transferred onto a glass slide (MATSUNAMI #S2441), gently covered with a glass coverslip, and observed using BZ-X710 (Keyence) with 20x and 40x objective lenses.

#### Isolation and analysis of barcoded colonies

After the barcode-specific Zeocin-resistance marker activation in the barcoded cell population, barcodes of colonies from test and control conditions were analyzed by Sanger sequencing. For each colony, the barcode region was amplified by PCR in 20 µL volume, composed of 1 µL of cell suspension, 0.5 µL each of 10 µM forward primer KI#403 and 10 µM reverse primer KI#404, 0.2 µL of Phusion High-Fidelity DNA Polymerase (NEB #M0530L), 4 µL of Phusion HF Buffer (NEB #B0518S), and 0.4 µL of 10 mM dNTPs (Takara #4030), with the following thermal cycle condition: 98°C for 30 s, 30 cycles of 98°C for 10 s, 54°C for 20 s, and 72°C for 30 s, and then 72°C for 5 min for the final extension. The PCR products were analyzed with 2% agarose gel and transferred into 96-well PCR plate wells for clean-up with 20 µL of AMPure XP beads (Beckman Coulter #A63881) according to the manufacturer’s protocol. The Sanger sequencing was performed using a sequencing primer KI#403. The Sanger sequencing trace was analyzed using PySanger (https://github.com/ponnhide/PySanger).

### High-throughput sequencing and data analysis

#### High-throughput sequencing

The sequencing libraries were mixed with 20–30% of PhiX spike-in DNA control (Illumina #FC-110-3001) for better cluster generation on the flow cell and sequenced by Illumina MiSeq (MiSeq v3 150-cycles kit #MS-102-3001 or 300-cycles kit #MS-102-3003), HiSeq2500 (TruSeq rapid SBS kit v2 #FC-402-4022). Base calling was performed with bcl2fastq2 (version v2.20.0) to generate FASTQ files. The sequencing condition for each library and the NCBI Sequence Read Archive’s ID to each raw FASTQ file can be found in Table S3.

#### Barcode identification

##### Mammalian CloneSelect C→T and yeast and bacterial CloneSelect

Sequencing reads were aligned to the constant sequences of the designed library structure using NCBI BLAST+ (version 2.6.0) (*67*) with the blastn-short option to identify sample indices for demultiplexing and CloneSelect barcode sequences. To generate the barcode allowlist for the mammalian CloneSelect C→T mini-pool experiments, barcode sequences commonly identified in both the plasmid DNA library and genomic DNA library were first obtained. Their sequencing errors were then corrected using starcode (version 1.4) (https://github.com/gui11aume/starcode) with the maximum Levenshtein distance threshold of 4 to merge minor and major barcodes. The barcode counts of each sample were normalized by the total barcode count. To estimate barcode frequencies for a given cell or DNA pool sample, the barcode frequencies were averaged across replicates if any. The barcode sequence and frequency information obtained in this study can be found in Table S1. Codes are available at https://github.com/yachielab/CloneSelect_v1/tree/main/Barcode_identification/CloneSelect.

##### scCloneSelect

To identify uptag and dntag barcodes of the barcoded cells, we first demultiplexed the sequencing reads and used cutadapt v4.1 (https://github.com/marcelm/cutadapt/) to extract uptag and dntag sequences sandwiched between their 20-bp upstream and downstream constant sequences. The identified uptags and dntags were filtered with the Q-score threshold of 30, clustered, and further filtered with their lengths of 17-bp for uptags and 30-bp for dntags using bartender-1.1 (https://github.com/LaoZZZZZ/bartender-1.1) (*68*). For the construction of the uptag-dntag combination reference database, uptags and dntags found in the same reads were first paired. We discarded redundant uptag-dntag pairs whose uptag or dntag were found in more abundant pairs. For the mapping of dntag to the uptag-dntag database, we used symspellpy v6.7 (https://github.com/mammothb/symspellpy) to find one with the shortest edit distance. In cases where multiple dntags in the database were found to be mapped by a query with the same edit distances, the dntag with the highest frequency in the database was chosen. To analyze uptag frequencies in a cell population obtained after gRNA-dependent barcode-specific EGFP reporter activation followed by flow cytometry cell sorting, we mapped the sequencing reads to the uptag-dntag database and obtained read counts for uptags recorded in the database using bartender-1.1 and symspellpy v6.7. Conflicts by the redundant hits to the database were resolved in the same way as those for dntags described above. Codes are available at https://github.com/yachielab/CloneSelect_v1/tree/main/Barcode_identification/scCloneSelect.

#### Drop-seq data analysis

After sample demultiplexing, FASTQ files were processed with Dropseq Tools v2.5.1 (https://github.com/broadinstitute/Drop-seq) for base quality filtering, adapter trimming, and Cell ID and UMI extraction. Picard v2.18.14 (https://github.com/broadinstitute/picard) was used to convert BAM files to FASTQ files for the proceeding step. The filtered reads were aligned using STAR v2.7 (https://github.com/alexdobin/STAR) (*69*) using the mm10 reference genome. The differential gene expression and clustering analysis were performed using Seurat v3 (https://github.com/satijalab/seurat) (*70*). After filtering cells with the thresholds of Feature_RNA > 200 & nFeature_RNA < 2500 & percent.mt < 5, their gene expression profiles were normalized before clustering with the Seurat::sctransform function. The dntags to be analyzed in this study were first determined in the first Drop-seq run based on their cumulative read count distribution of Cell IDs with the threshold determined by the knee point using the Python package kneed v0.8.1 (https://github.com/arvkevi/kneed). However, to map dntag distribution to the single-cell transcriptome data with high sensitivity, we also sequenced the dntag-reamplified library and identified the association of Cell IDs and dntags in the dntag-uptag combination reference database by the method described above for the scCloneSelect uptag and dntag identification. In cases where multiple dntags were found for a single Cell ID, the dntag with the most abundant UMI count was chosen. Codes are available at https://github.com/yachielab/CloneSelect_v1/tree/main/Drop-seq.

#### Mutational spectra analysis

The amplicon sequencing reads obtained to analyze mutational patterns at the CAN1 target site conferred by each of the Cas9-based genome editing enzymes was processed by a pipeline described previously (*39*). Codes specific to this study are available at https://github.com/yachielab/CloneSelect_v1/tree/main/Mutational_Spectra_Analysis.

### Statistical analysis

The statistical tests were performed using R (version 4.2.0). The details of each statistical test can be found in the corresponding figure legend. The statistical methods used in this study and *P*-values are also listed in Table S4.

### Data and code availability

High-throughput sequencing data generated in this study are available at the NCBI Sequence Read Archive (PRJNA901977). All the codes used in this study are available at https://github.com/yachielab/CloneSelect_v1.

## Supplementary Materials for

**Fig. S1.**
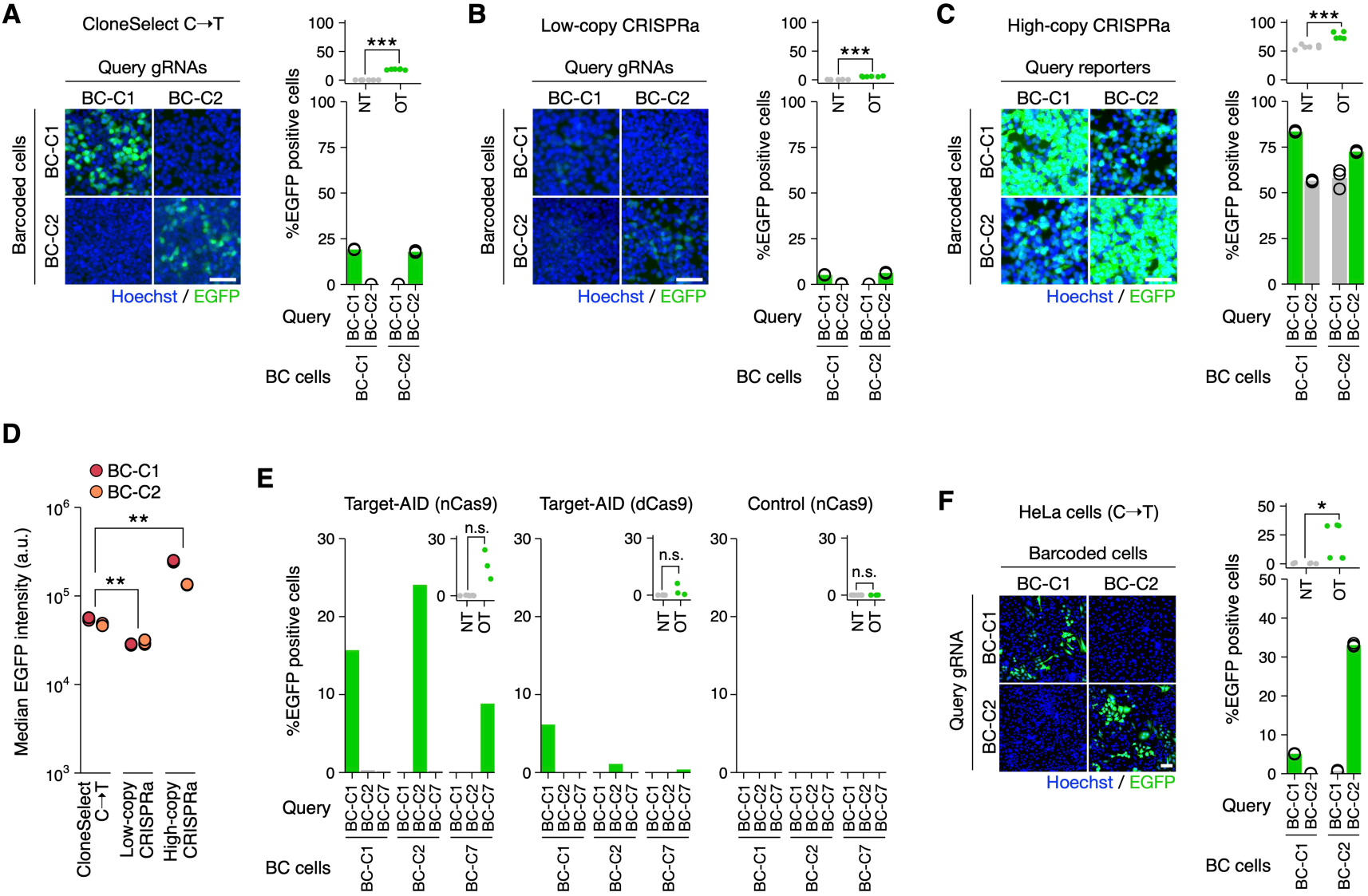
Supplementary data for developing CloneSelect C→T. (**A**–**C**) Barcode-specific gRNA-dependent activation of EGFP reporters for two barcoded HEK293T strains established for each of CloneSelect C→T (**A**), low-copy CRISPRa (**B**), and high-copy CRISPRa (**C**) (n=3). Scale bar, 50 µm. Welch’s t-test was performed to compare on-target (OT) and non-target (NT) activations. (**D**) Median EGFP intensities of genome editing-activated EGFP positive cells (n=3). The Mann-Whitney U test was performed to compare two groups. (**E**) Comparison of Target-AID variants and a nCas9 (D10A) control in the CloneSelect C→T reporter activation for the same set of barcode-gRNA pairs (n=1). Welch’s t-test was performed to compare OT and NT activations. (**F**) Reporter activation in HeLa cells by CloneSelect C→T (n=3). Welch’s t-test was performed to compare OT and NT activations. Scale bar, 80 µm. \**P* < 0.05; \*\**P* < 0.01; \*\*\**P* < 0.001.

**Fig. S2.**
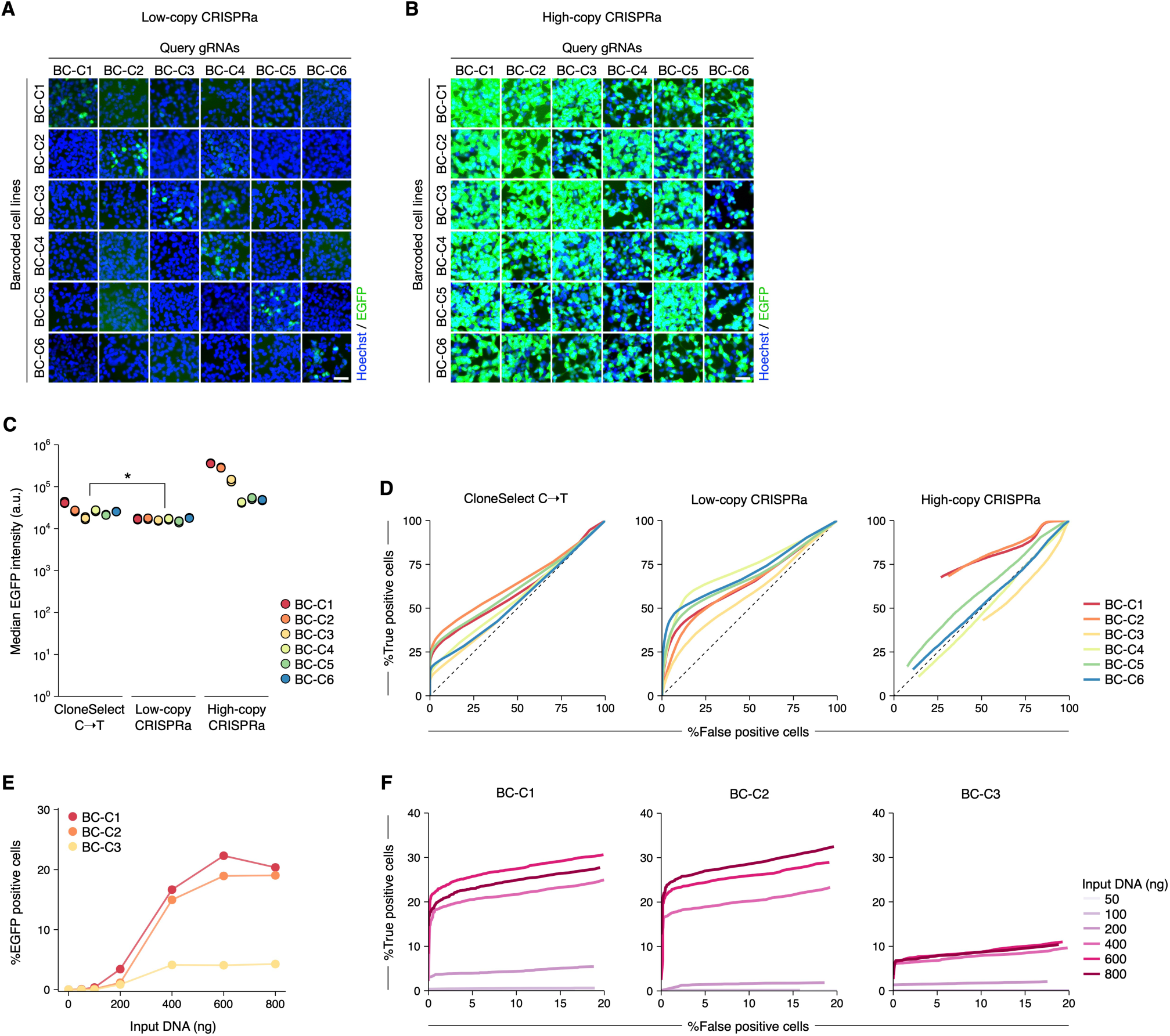
Supplementary data for comparing CloneSelect C→T with CRISPRa-based circuits. (**A** and **B**) Barcode-specific gRNA-dependent reporter activation of six barcoded cell lines prepared for each of low-copy CRISPRa and high-copy CRISPRa. Scale bar, 50 µm. (**C**) Median EGFP intensities of genome editing-activated EGFP positive cells (n=3). (**D**) ROC curves along varying reporter intensity thresholds for target barcoded cells. CloneSelect C→T, low-copy CRISPRa, and high-copy CRISPRa were examined for the common set of six barcodes. The Mann-Whitney U test was performed to compare two groups (\**P* < 0.05; \*\**P* < 0.01; \*\*\**P* < 0.001). (**E**) Frequencies of CloneSelect C→T reporter-activated cells obtained by transfection of different DNA amounts of barcode-targeting genome editing reagents. (**F**) ROC curve for each input DNA amount along varying reporter intensity thresholds for target barcoded cells.

**Fig. S3.**
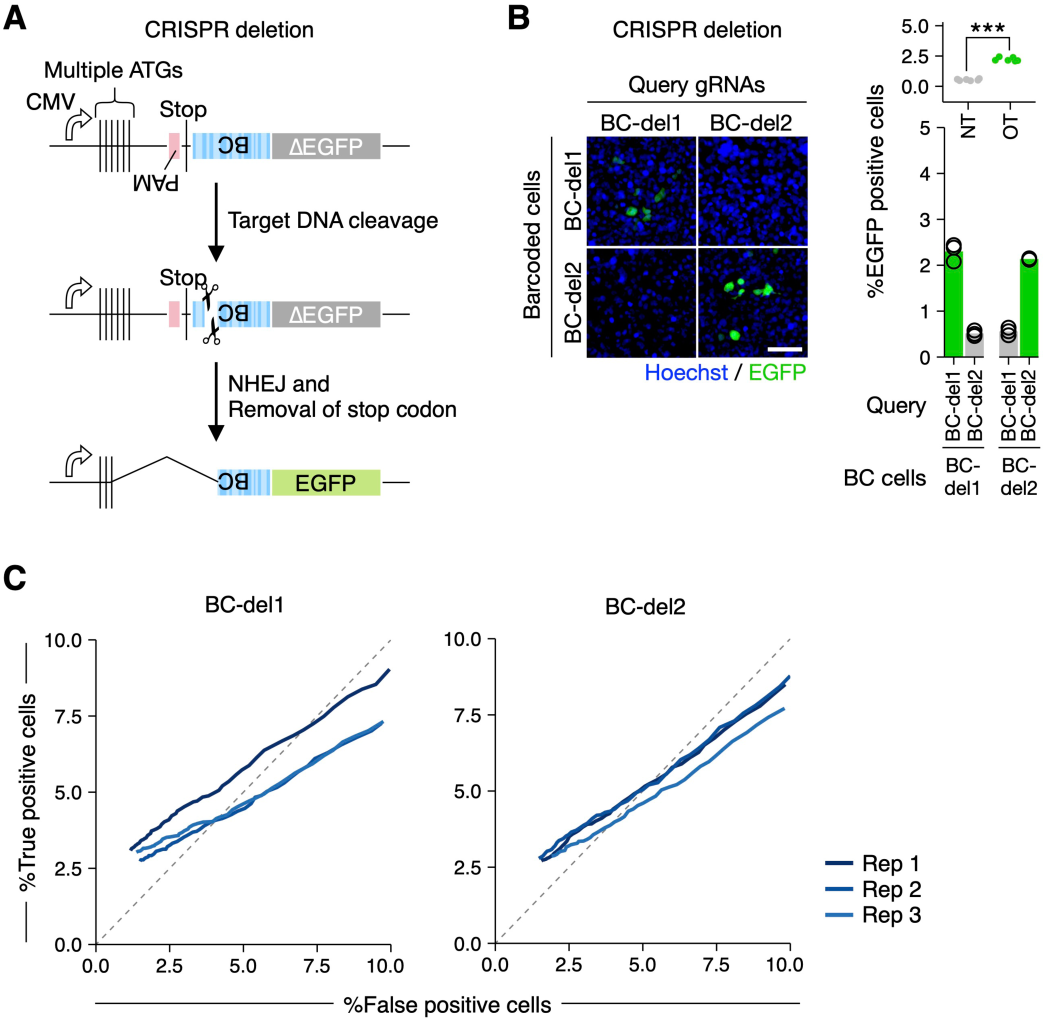
DSB-based circuit. (**A**) Barcode-specific gRNA-dependent reporter activation circuit using wild-type Cas9. (**B**) Barcode-specific activation of the deletion-based reporter prepared for two barcodes (BC-del1 and BC-del2) in HEK293T cells (n=3). Welch’s t-test was performed to compare on-target (OT) and non-target (NT) activations (\**P* < 0.05; \*\**P* < 0.01; \*\*\**P* < 0.001). Scale bar, 50 µm. (**C**) ROC curves along varying reporter intensity thresholds for target barcoded cells (n=3).

**Fig. S4.**
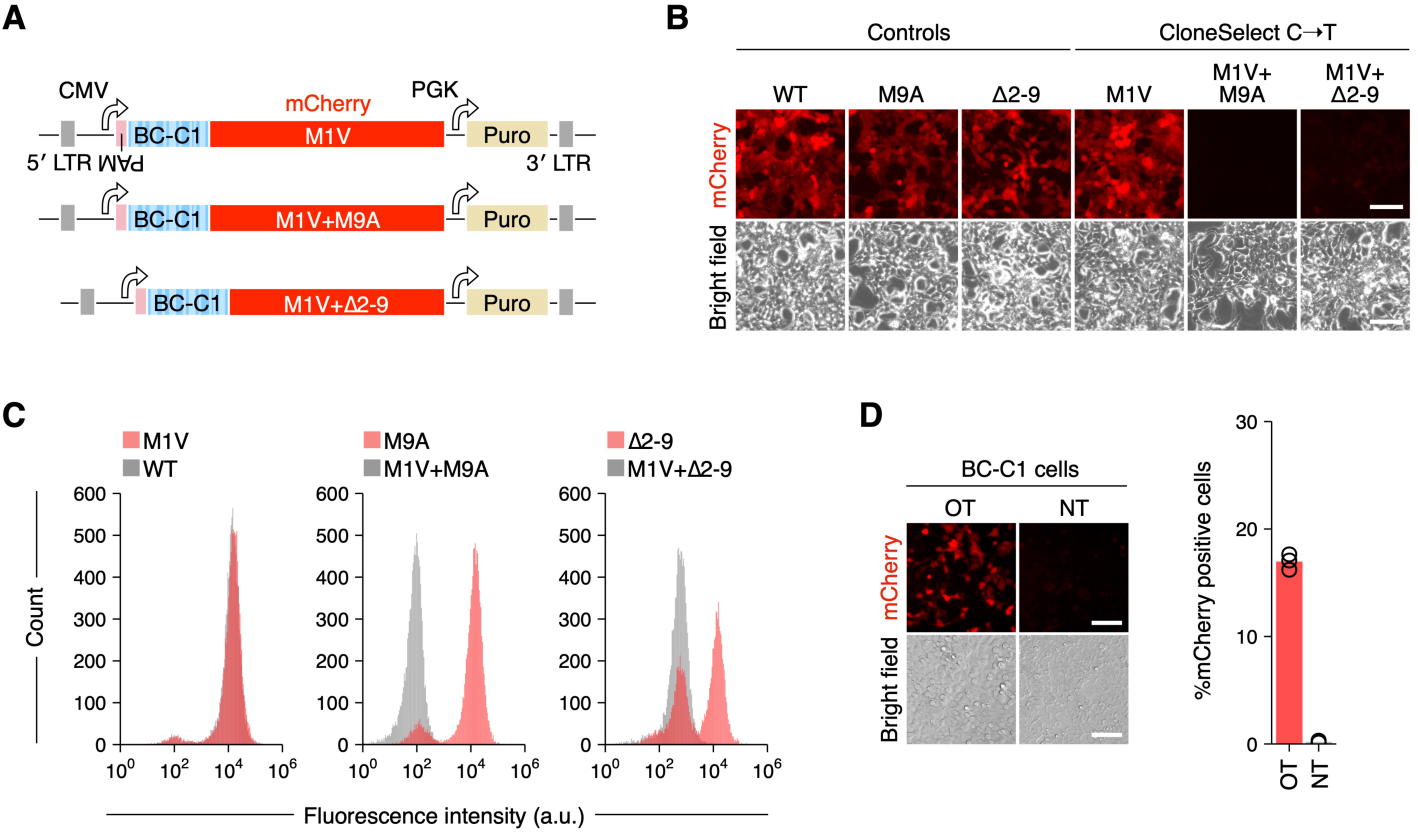
mCherry reporter for CloneSelect C→T. (**A**) Different mCherry reporter variants tested to establish CloneSelect C→T. (**B** and **C**) mCherry expression from the different reporter variants with the first codon as GTG or ATG. Scale bar, 50 µm. (**D**) Activation of the M1V (GTG)+Δ2-9 mutant reporter with OT and NT gRNAs (n=3). Scale bar, 100 µm.

**Fig. S5.**
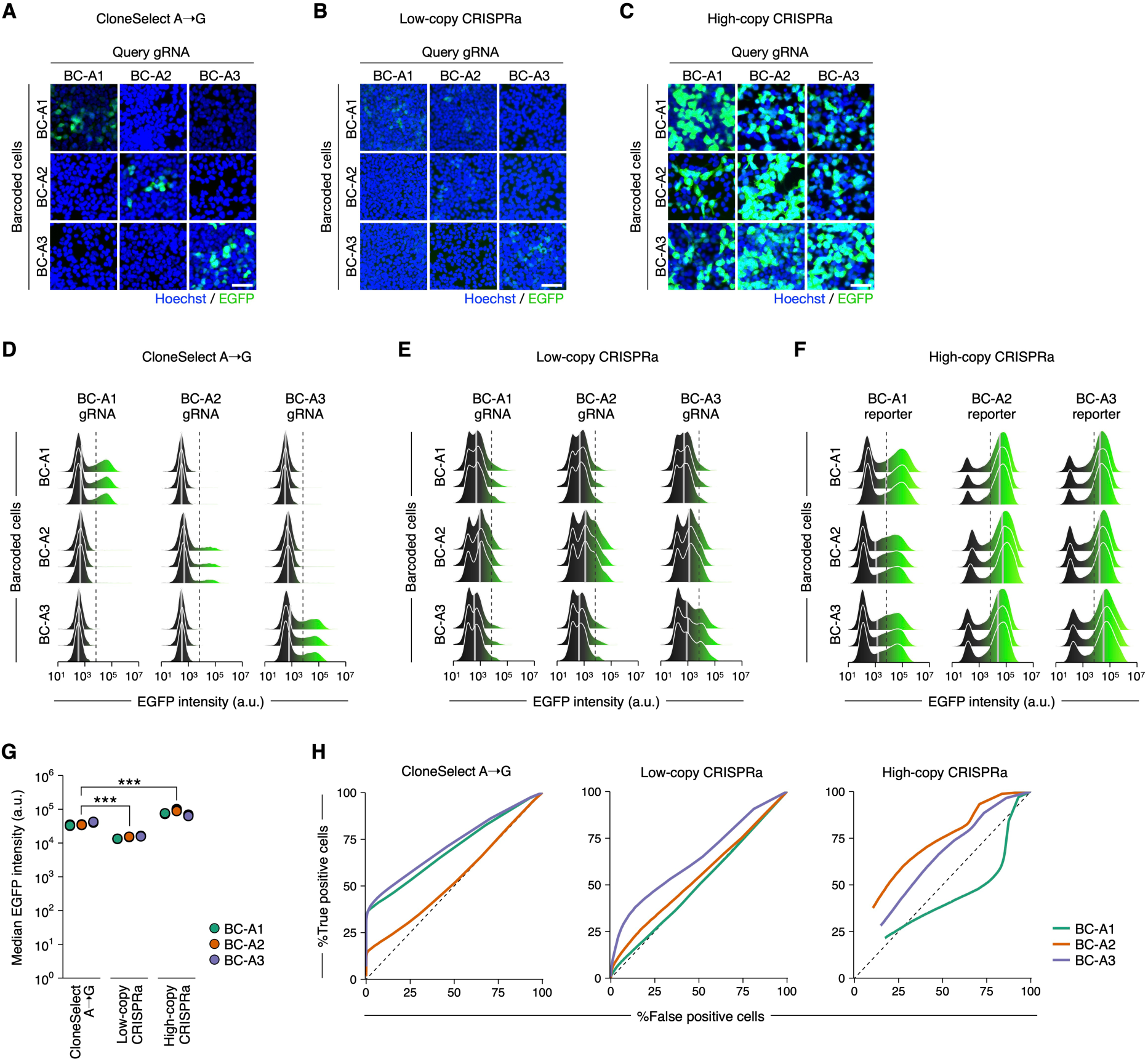
Supplementary data for comparing CloneSelect A→G with CRISPRa-based circuits. (**A**–**C**) Barcode-specific gRNA-dependent reporter activation of three barcoded cell lines prepared for each of CloneSelect A→G, low-copy CRISPRa, and high-copy CRISPRa. Scale bar, 50 µm. (**D**–**F**) Flow cytometry analysis of single-cell EGFP activation levels. (**G**) Median EGFP intensities of genome editing-activated EGFP positive cells (n=3). The Mann-Whitney U test was performed to compare two groups (\**P* < 0.05; \*\**P* < 0.01; \*\*\**P* < 0.001). (**H**) ROC curves along varying reporter intensity thresholds for target barcoded cells (n=3).

**Fig. S6.**
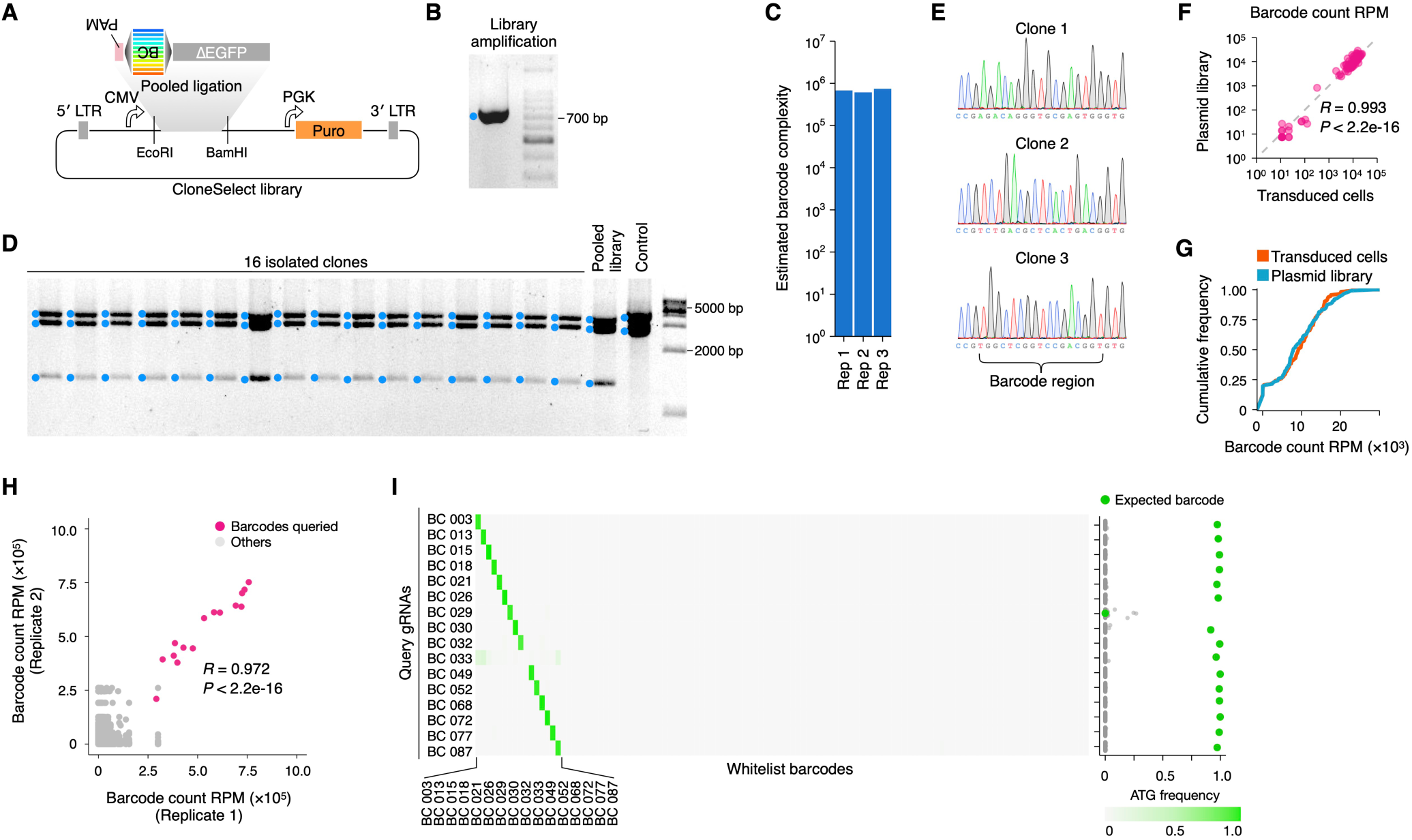
Supplementary data for isolating barcoded cells using CloneSelect C→T. (**A**) Schematic diagram for the barcode library construction. The ΔEGFP fragment (no start codon) was amplified by PCR using pooled forward primers encoding the PAM followed by semi-random barcode sequences encoding GTG and a common reverse primer, and enzymatically digested and ligated to the lentiviral backbone. (**B**) The insert PCR fragments. (**C**) Estimated complexities of the generated plasmid pools in triplicate. (**D**) Library QC by colony isolation and restriction digest by BsrGI, ClaI and PvuI. (**E**) Sanger sequencing of the barcode region of the colony isolates. (**F** and **G**) Barcode distribution in the lentiviral plasmid DNA pool and that in the cell population transduced using the same plasmid DNA pool. (**H**) Barcode distribution in the EGFP-positive cell population obtained by cell sorting after barcode-specific activation by a target gRNA. The results of the 16 independent samples were combined after read count normalization for each sample. (**I**) Frequency of the GTG→ATG mutation observed for each barcode after sorting of the reporter positive cells. Each row represents the GTG→ATG mutation frequency profile obtained for each target isolation attempt.

**Fig. S7.**
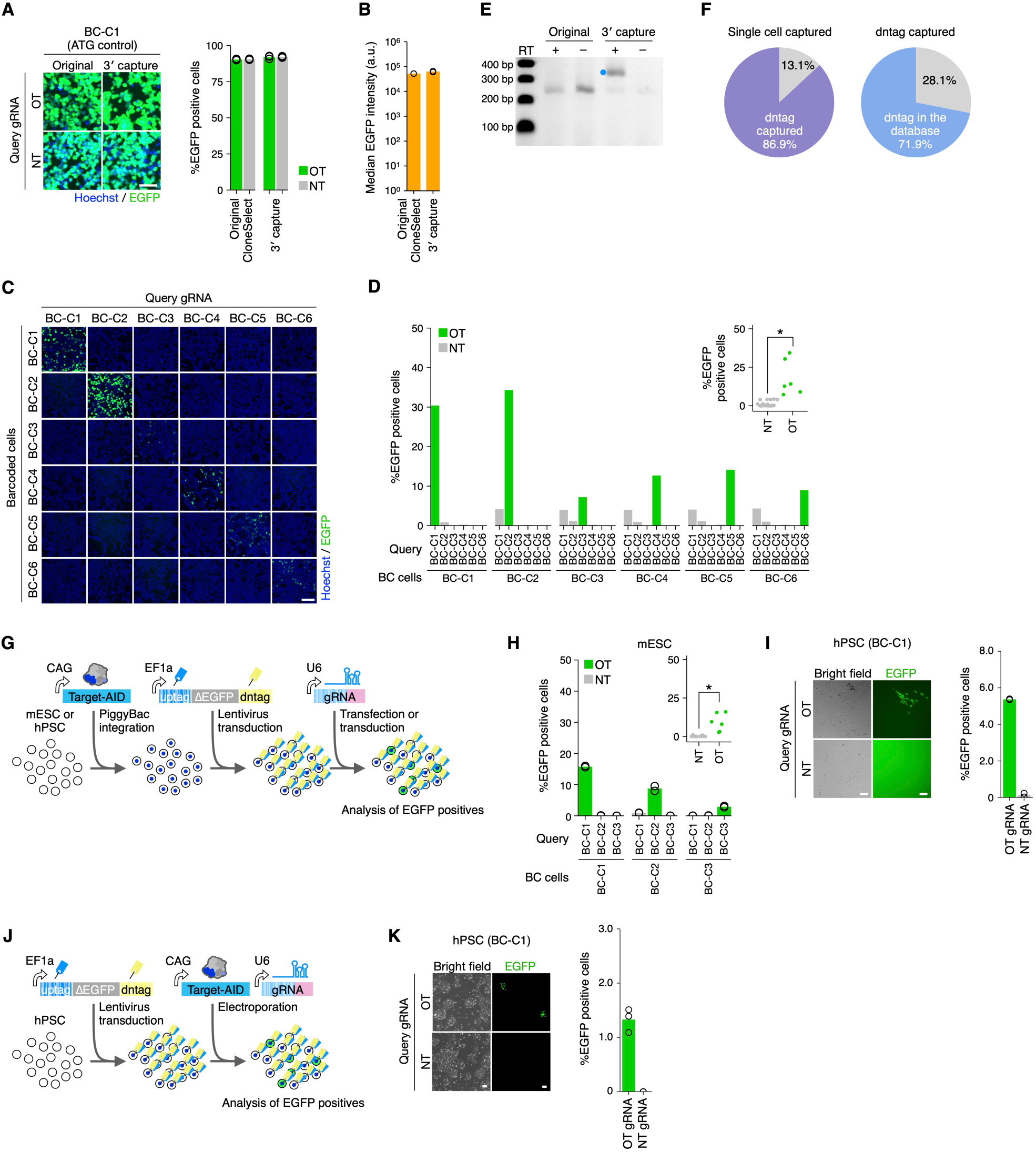
Supplementary data for developing scCloneSelect. (**A**) EGFP-positive control expressions for the original CloneSelect C→T and scCloneSelect in HEK293T cells with the same genome editing conditions tested for the respective reporters (n=3). Scale bar, 50 µm. (**B**) Median EGFP intensities of base editing-activated EGFP positive cells (n=3). (**C** and **D**) Barcode-specific gRNA-dependent reporter activation of six barcoded cell lines by scCloneSelect (n=1). Welch’s t-test was performed to compare on-target (OT) and non-target (NT) activations (\**P* < 0.05; \*\**P* < 0.01; \*\*\**P* < 0.001). Scale bar, 50 µm. (**E**) RT-PCR of the scCloneSelect dntags in HEK293T. (**F**) Fraction of mESC single-cell transcriptome profiles (Drop-seq) that contained dntags and fraction of dntags reported in the uptag-dntag combination reference database. (**G**) Schematic representation of a scCloneSelect reporter activation assay where Target-AID was stably introduced to the cell population prior to barcoding and gRNA-dependent reporter activation. (**H** and **I**) gRNA-dependent reporter activation of target barcoded mESCs and CA1 hPSCs by scCloneSelect (n=2). Target-AID was stably integrated prior to the barcoding. Targeting gRNAs were delivered by transfection. Welch’s t-test was performed to compare OT and NT activations (\**P* < 0.05; \*\**P* < 0.01; \*\*\**P* < 0.001). Scale bar, 100 µm. (**J**) Schematic representation of a scCloneSelect reporter activation assay where the target gRNA and Target-AID were electroporated together to the barcoded cell population. (**K**) gRNA-dependent reporter activation of barcoded H1 hPSCs by scCloneSelect (n=2). Targeting gRNA and Target-AID were electroporated together. Scale bar, 100 µm.

**Fig. S8.**
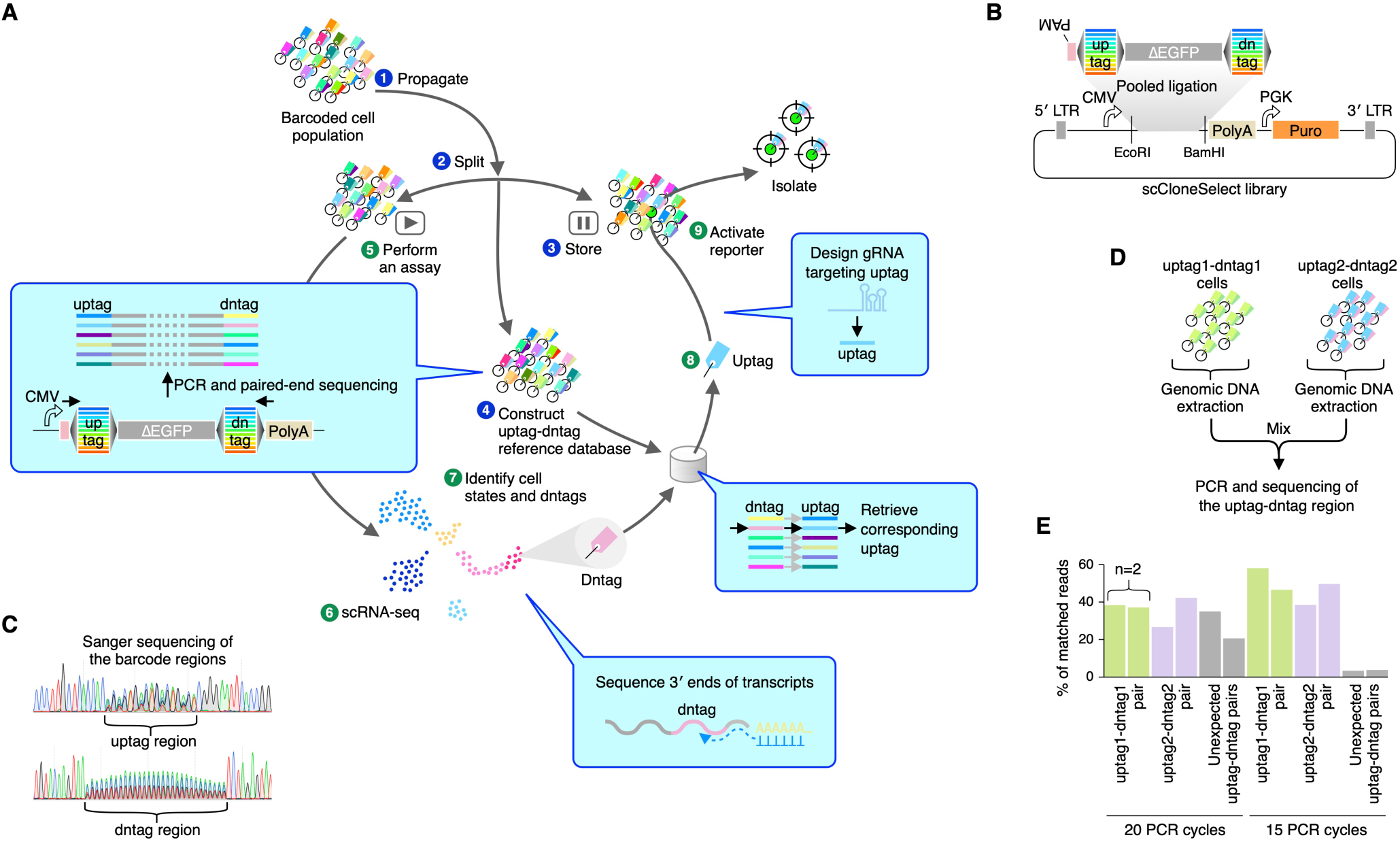
The gene expression profile cytometry cell sorting concept. (**A**) Schematic diagram of a scCloneSelect workflow. (**B**) Preparation of barcode library for scCloneSelect. (**C**) Sanger sequencing results of the uptag and dntag regions of the constructed plasmid pool. (**D**) Mixing of genomic DNA samples from two barcoded cell lines to optimize the PCR protocol to create a library for identifying the uptag-dntag combination reference database. (**E**) Fraction of unexpected chimeric PCR recombination products in the sequencing reads.

**Fig. S9.**
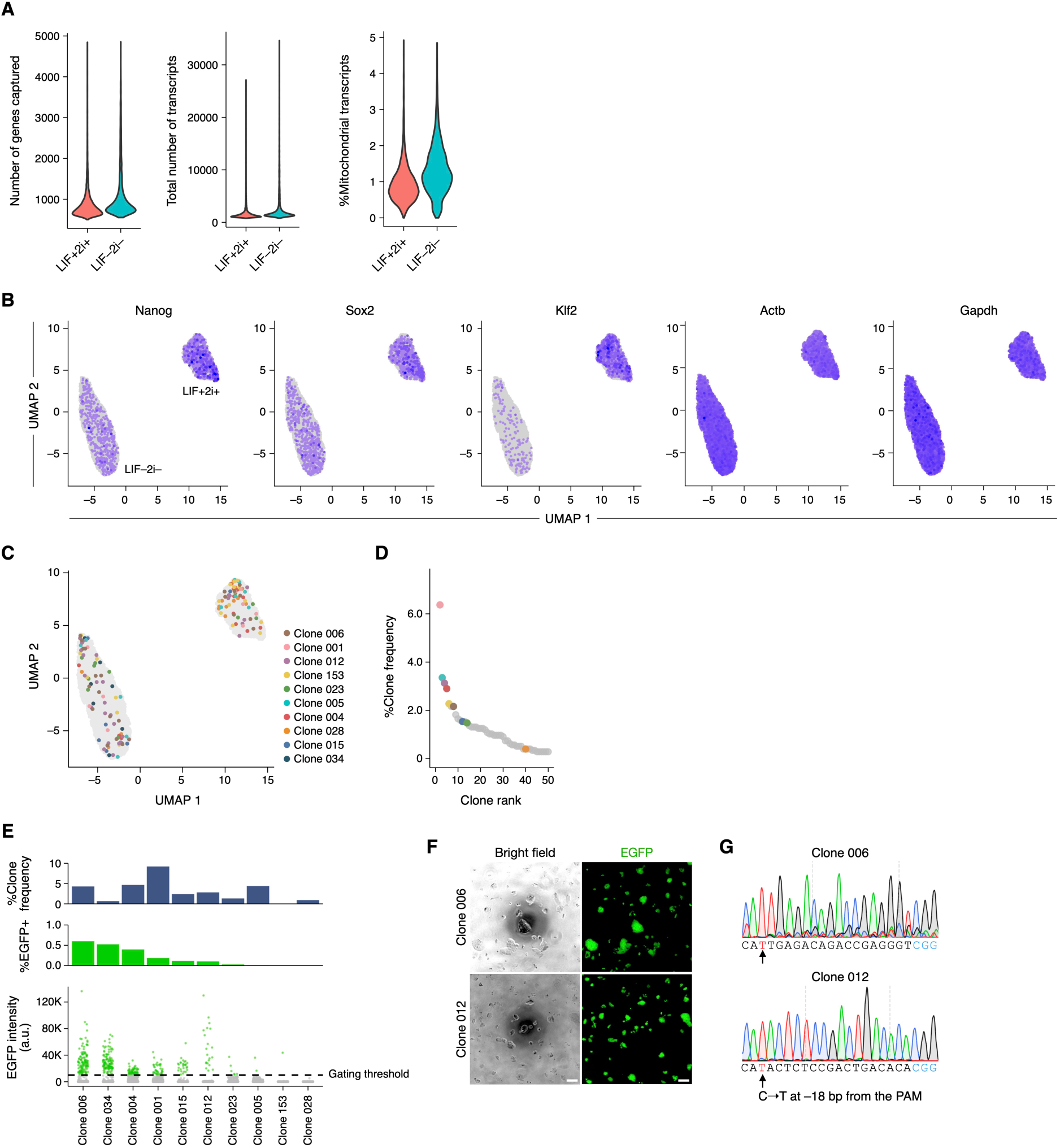
Supplementary data for isolating barcoded cells identified in scRNA-seq data. (**A**) QC of scRNA-seq datasets obtained for the barcoded mESC population cultured with LIF and 2i and that cultured without LIF or 2i. (**B**) Single-cell expression patterns of key genes. (**C**) Distribution of cells in a two-dimensional UMAP space for all the clones targeted for isolation. (**D**) Abundances of barcoded cell clones in the mESC population. The data was generated based on dntags identified in the scRNA-seq dataset with no reamplification of the dntag reads. (**E**) gRNA-dependent activation of the reporter for each of the target clones in the initial mESC population. (**F**) Culturing of single isolated cells. Scale bar, 100 µm. (**G**) Sanger sequencing of the barcodes for the isolated target clones expanded after clone labeling and single-cell isolation.

**Fig. S10.**
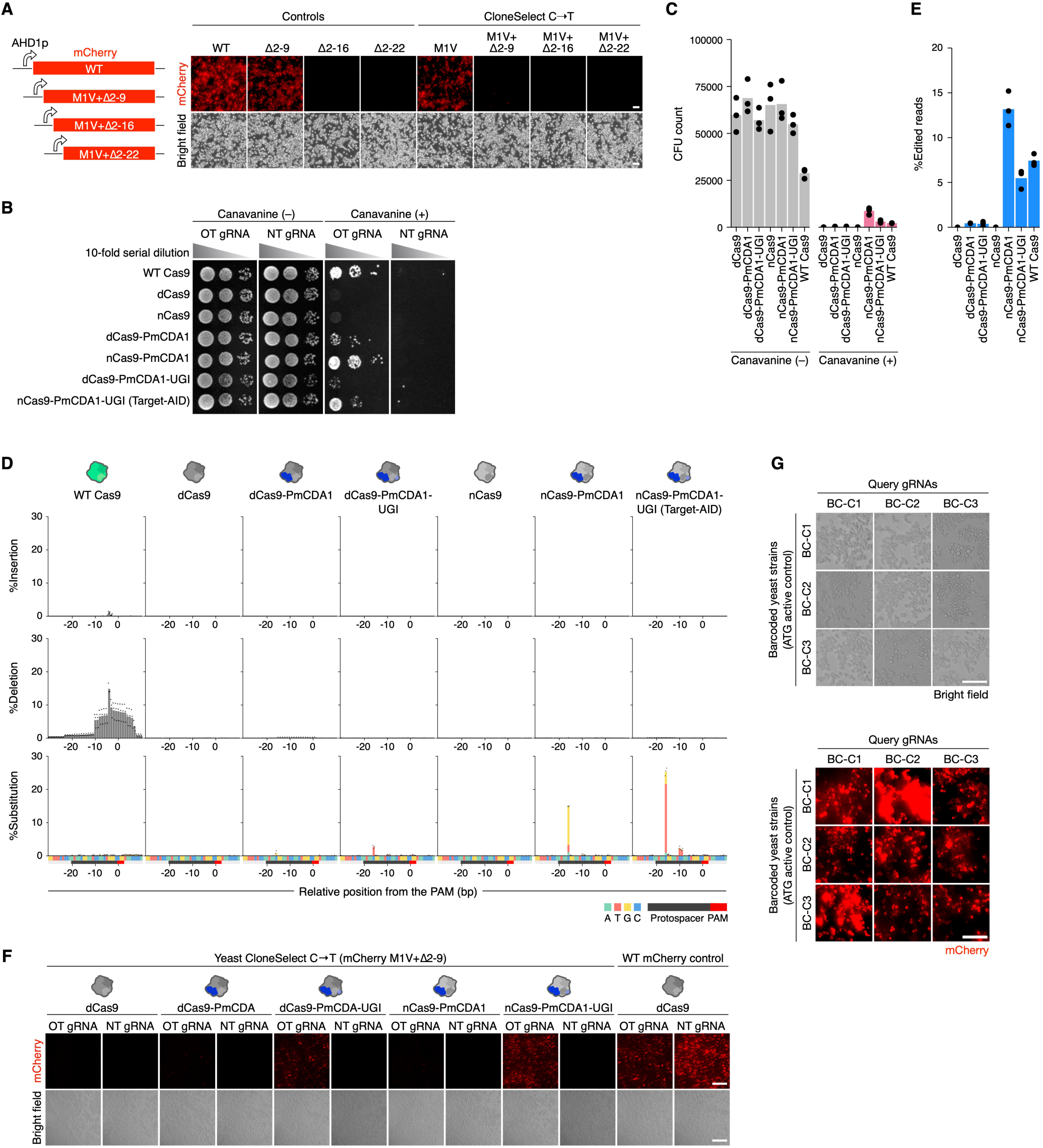
Supplementary data for developing yeast CloneSelect. (**A**) Different mCherry reporter variants tested to establish CloneSelect C→T. The different reporter variants were tested with the first codon as GTG or ATG. Scale bar, 100 µm. (**B**) Canavanine resistance assays for different CRISPR genome editing enzymes with a gRNA targeting CAN1 gene and a control NT gRNA. For each experiment, cell concentration was normalized to 1.0 OD_595 nm_ and serially diluted with 10-fold increments for spotting. (**C**) Estimated CFU counts for the same assay in (**B**). (**D**) Genome editing outcomes observed by amplicon sequencing. Frequencies of mutation patterns observed across the target sequence region are shown for the same assay in (**B**). (**E**) Genome editing frequencies at the target CAN1 locus estimated by amplicon sequencing for the different enzymes. (**F**) Activation of the mCherry M1V (GTG)+Δ2-9 mutant reporter by OT and NT gRNAs. Scale bar, 200 µm. (**G**) mCherry-positive control expressions for yeast CloneSelect. Yeast cells having the positive control reporters with three different barcodes (BC-C1, BC-C2, and BC-C3) were each treated by Target-AID and three different targeting gRNAs. Scale bar, 25 µm.

**Fig. S11.**
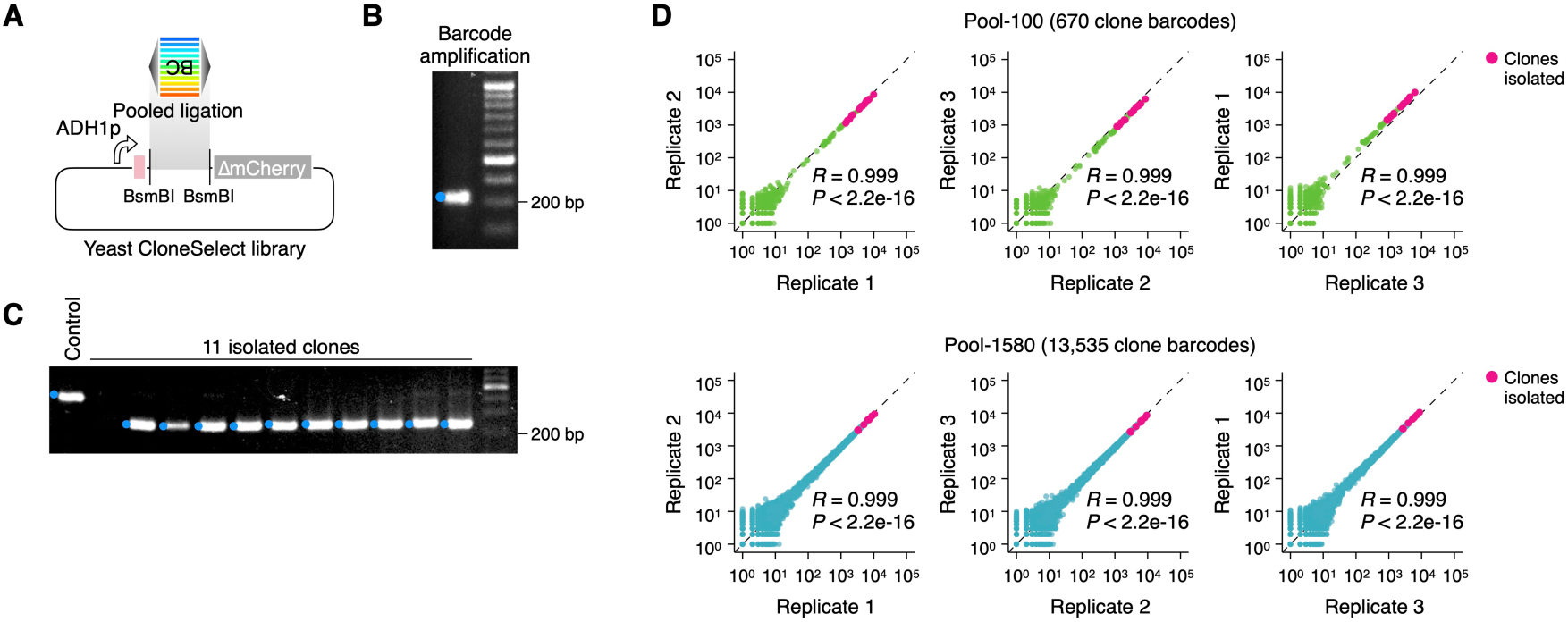
Preparation of yeast CloneSelect library. (**A**) Schematic diagram for the barcode library construction. The barcode fragment pool was prepared by PCR using a common primer pair amplifying a template DNA pool encoding the PAM, WSNS semi-random repeat, and the mutated start codon GTG. The PCR product was digested and ligated to a backbone plasmid. (**B**) The insert PCR fragments (Pool-100). (**C**) Library QC by colony isolation and PCR amplification of the barcode insert (Pool-100). (**D**) Barcode abundance distribution (read per million) in the constructed barcode library pool. The sequencing library was prepared and analyzed in triplicate (n=3).

**Fig. S12.**
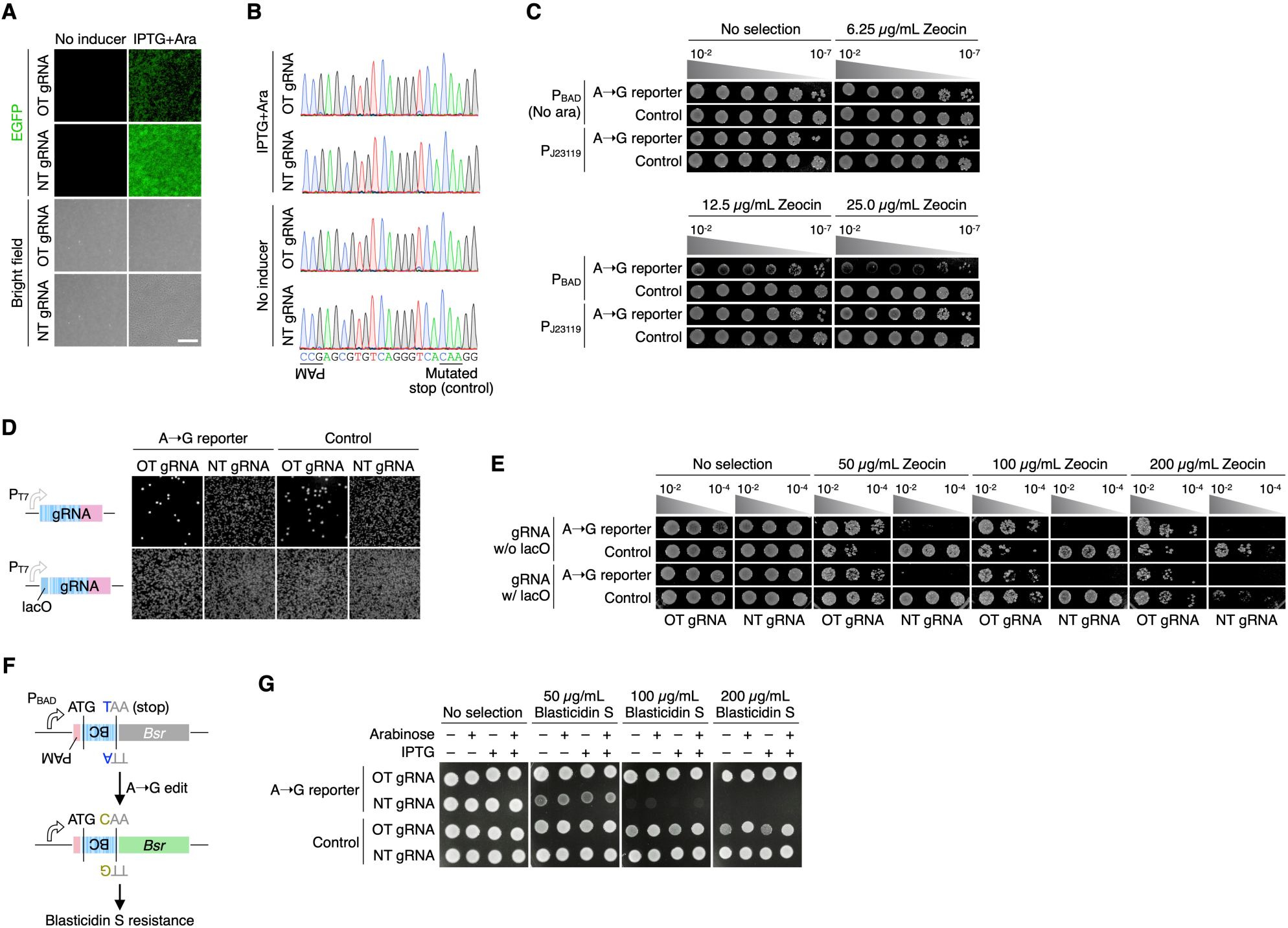
Supplementary data for developing bacterial CloneSelect. (**A**) Activities of the positive control EGFP reporter. ABE and gRNA expression were controlled by an IPTG-inducible promoter, and the EGFP reporter expression was controlled by an arabinose-inducible promoter. (**B**) Base editing outcomes of the positive control reporters analyzed by Sanger sequencing. (**C**) Testing of Zeocin resistances conferred by two promoters expressing a Zeocin resistance gene with and without the upstream stop codon to block the selective marker translation. Each cell sample concentration was first adjusted to 0.1 OD_595 nm_ and serially diluted with 10-fold increments for spotting 5 µL. (**D**) Testing of cell viability under a non-selective condition for a constitutively active T7 promoter and the IPTG-inducible promoter to express the gRNA. OT and NT gRNAs were tested for the gRNA-dependent EGFP reporter and the positive control EGFP reporter. ABE was expressed under the IPTG-inducible promoter without IPTG provided. (**E**) gRNA-dependent Zeocin resistance reporter activation tested for the IPTG-inducible promoters with and without IPTG. (**F**) Bacterial CloneSelect using the Blasticidin resistance gene. Each cell sample concentration was first adjusted to 0.1 OD_595 nm_ and serially diluted with 10-fold increments for spotting 5 µL. (**G**) gRNA-dependent Blasticidin-resistance reporter activation tested for different inducer conditions and different Blasticidin concentrations. Each cell sample concentration was adjusted to 0.1 OD_595 nm_ for spotting 5 µL.

**Fig. S13.**
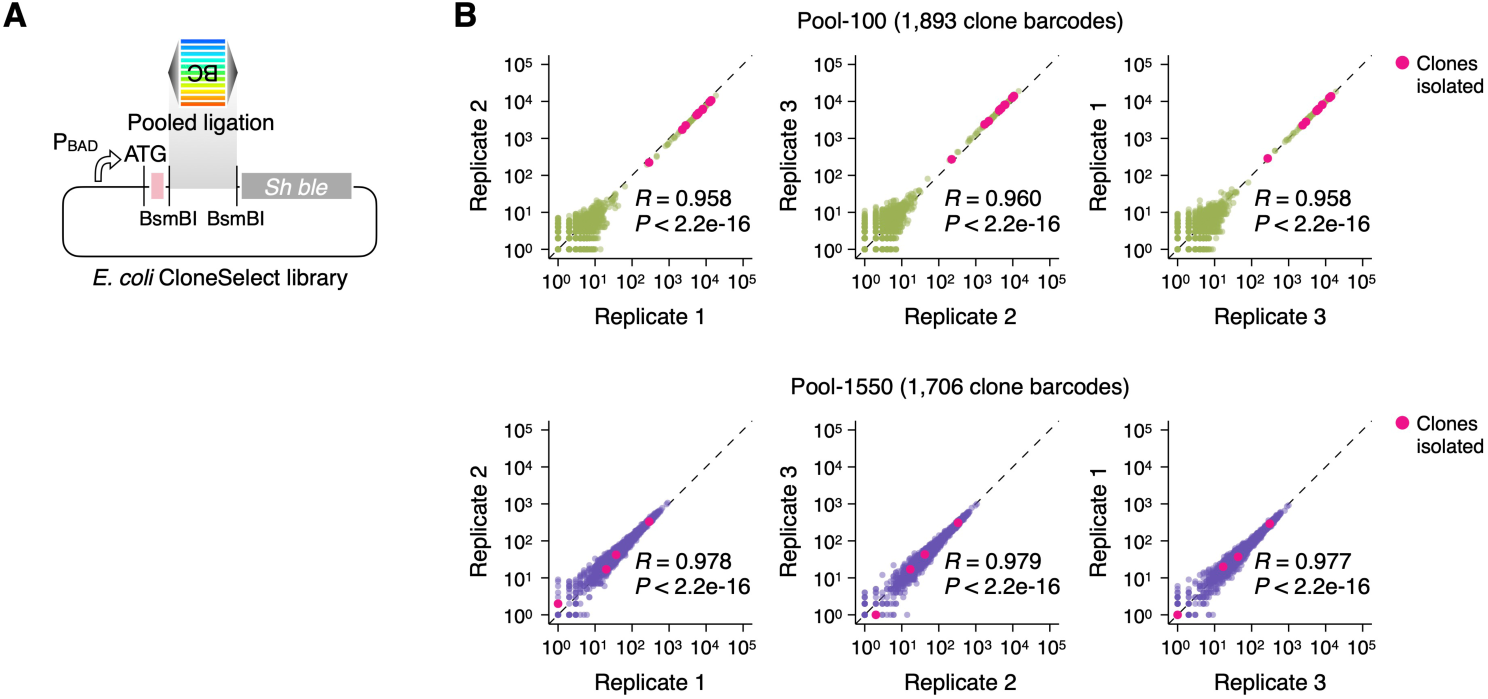
Preparation of *E. coli* CloneSelect library. (**A**) Schematic diagram for the barcode library construction. The barcode fragment pool was prepared by PCR using a common primer pair amplifying a template DNA pool encoding the PAM, VNN repeat sequence, and the start codon TAA. The PCR product was digested and ligated to a backbone plasmid. (**B**) Barcode abundance distribution (read per million) in the constructed barcode library pool. The sequencing library was prepared and analyzed in triplicate (n=3).

**Fig. S14.**
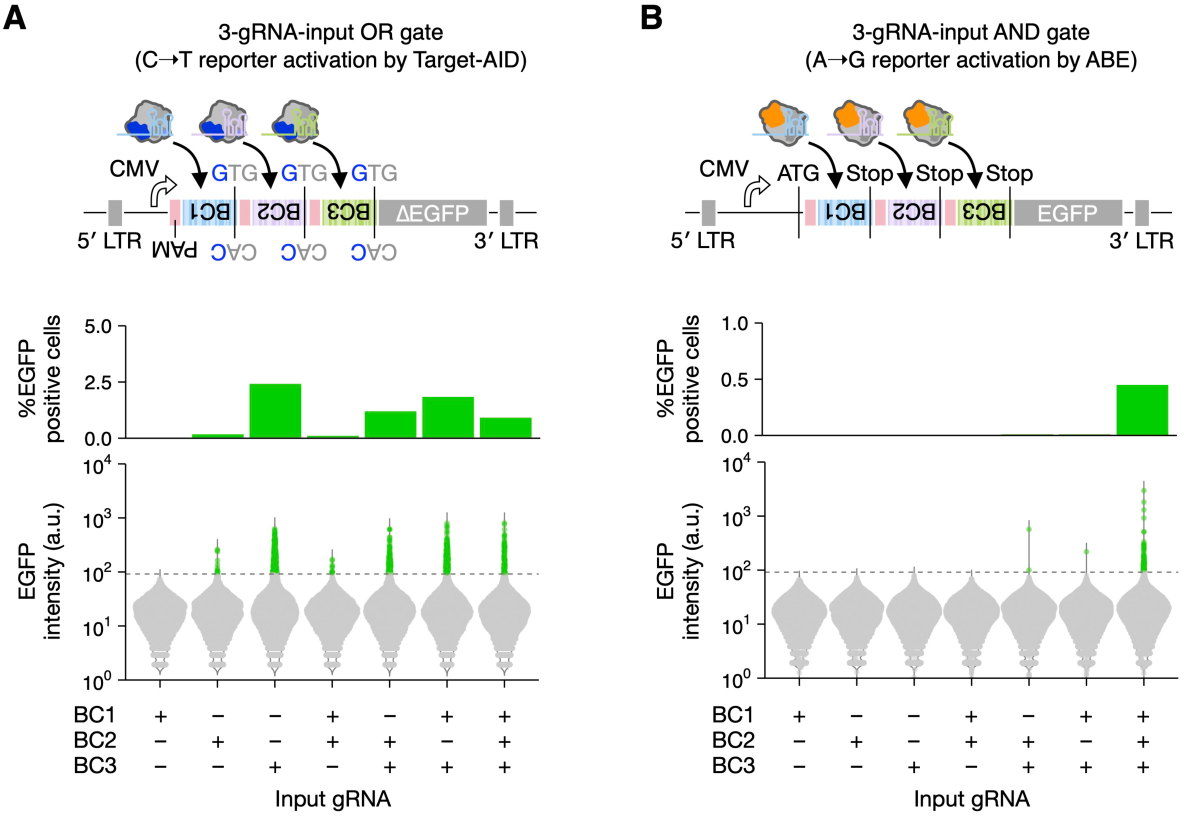
Multiple gRNA-input logic gates. (**A**) Three-gRNA-input OR gate with CloneSelect C→T that is designed to confer the EGFP reporter expression by any of the three barcode-specific gRNA-dependent GTG→ATG mutations. (**B**) Three-gRNA-input AND gate with CloneSelect A→G that is designed to confer the EGFP reporter expression when all three barcode-specific gRNA-dependent TAA→CAA mutations are provided.

